# Machine-assisted systematic review and meta-analysis of citrus flavonoids in streptozotocin-induced painful diabetic neuropathy

**DOI:** 10.64898/2026.09.27.750067

**Authors:** Shreya Jindal, Jayashrii Muthaiyan, Sean Smith, Nadia Soliman, Gillian L. Currie

**Affiliations:** University of Edinburgh; University College London

## Abstract

**Background:** Neuropathic pain is a common and debilitating condition with persistently poor translation from preclinical research to effective clinical therapies. Preclinical systematic reviews and meta-analyses (SRMAs) can provide more robust estimates of intervention effects and help identify sources of bias and heterogeneity, but their use is limited by the time and effort required to synthesise rapidly expanding literatures.

Automation may help overcome these barriers.

**Methods:** We developed Neuropathic Pain - Systematic Online Living Evidence Summaries (NP-SOLES), a machine-assisted approach used in this study to support systematic identification of animal studies relevant to neuropathic pain. Using NP-SOLES, we conducted a pilot systematic review and meta-analysis of controlled animal studies assessing the effects of citrus flavonoids on thermal and mechanical pain-related outcomes in streptozotocin-induced models of painful diabetic neuropathy.

**Results:** In this pilot application, machine-assisted evidence synthesis supported reduced manual screening burden, illustrating the potential of such approaches to facilitate timely preclinical SRMAs. Fourteen studies met inclusion criteria for meta-analysis on thermal pain-related outcomes. Pooled estimates suggested a reduction in thermal pain-related behaviours following flavonoid treatment; however, substantial heterogeneity and widespread methodological limitations were observed. Nine studies were eligible for qualitative analysis on mechanical pain-related outcomes, which showed an overall reduction in pain-related behaviours following treatment but had similar methodological limitations. Reporting of key measures to reduce risk of bias, including randomisation and blinding, were consistently poor, and subgroup analyses were constrained by sparse data.

**Conclusions:** Available evidence from preclinical animal studies suggests a potential analgesic effect of citrus flavonoids on thermal and mechanical pain-related outcomes in diabetic neuropathy models, but confidence in this estimate is limited by significant risks of bias. More rigorously designed and transparently reported preclinical studies are required before clinical translation can be justified.

## Background

Neuropathic pain is a chronic and debilitating condition arising from damage or disease affecting the somatosensory system (Jensen *et al*., 2011), with substantial impacts on quality of life and limited effective treatment options (van Velzen, Dahan and Niesters,2020). The prevalence of neuropathic pain in the general population has been estimated to range from 3% to 17%, with painful diabetic neuropathy (PDN) representing a substantial contributor to this burden (Cavalli *et al*., 2019). This number is only expected to increase due to factors such as rising rates of diabetes mellitus and an ageing population (Colloca *et al*., 2017). PDN can present with pain that is spontaneous; typically described as electric shock-like, and evoked pain including allodynia and hyperalgesia (Finnerup, Kuner and Jensen, 2020). Current pharmacological treatments provide incomplete relief for many patients and are often associated with adverse effects (van Velzen, Dahan and Niesters, 2020).

Animal models have been central to efforts to understand PDN and to identify candidate analgesic interventions. Streptozotocin (STZ)-induced models are widely used to reproduce aspects of diabetic neuropathy, including alterations in nociceptive processing, and mechanical and thermal hypersensitivity (Fox *et al*., 1999). However, preclinical neuropathic pain research often has limitations of poor experimental design and quality (Jin, Moon and Calcutt, 2021). These factors contribute to inconsistent findings, reduced clinical translatability of research and may inflate estimates of treatment efficacy, helping to explain why apparently promising interventions frequently fail in clinical trials (Fisher *et al*., 2021).

PDN research over the years has highlighted a growing interest in the development of treatments from natural sources. An emerging candidate is citrus flavonoids; a class of phytochemical compounds found in citrus plants that are sustainable and low cost in production. They are reported to have analgesic effects in animal models and proposed biological actions, including antioxidant, anti-inflammatory and insulin-resistant properties (Wang *et al*., 2022). These mechanisms are biologically plausible given the roles of oxidative stress and inflammation in diabetic neuropathy (Vincent *et al*., 2011). However, evidence supporting flavonoid efficacy is dispersed across individual studies that vary widely in quality, model characteristics, and outcome assessment. Without systematic synthesis, it is difficult to assess the consistency and credibility of reported effects or to determine whether observed benefits are robust to methodological scrutiny.

Systematic reviews and meta-analyses allow estimation of overall treatment effects while interrogating heterogeneity and methodological bias. By aggregating results across studies, such reviews can help distinguish reliable treatment effects from findings driven by small sample sizes, selective reporting, or poor study design (Mulrow, 1994). Despite these advantages, preclinical systematic reviews remain relatively uncommon, in part because the volume and pace of publication make comprehensive manual screening and updating resource intensive (Ineichen *et al*., 2024).

In this study, we conducted a systematic review of controlled preclinical animal studies evaluating the effects of citrus flavonoids on thermal and mechanical pain-related outcomes in STZ-induced models of painful diabetic neuropathy. Thermal and mechanical outcomes were selected due to their clinical relevance in altered thermal sensitivity, and evoked allodynia and hyperalgesia respectively as well as being widely reported in STZ-induced models. A meta-analysis was conducted for thermal outcomes, however, was not conducted for mechanical outcomes where only descriptive analysis was carried out due to insufficient availability of data for quantitative synthesis according to our predetermined protocol. To support efficient identification of relevant studies within a large neuropathic pain literature, we developed and applied NP-SOLES, a machine-assisted screening approach as part of this review. In addition to estimating pooled treatment effects, we critically assessed study design and quality, and sources of bias to evaluate the strength and limitations of the available preclinical evidence.

## Methods

### Development of the NP-SOLES Machine Assisted Approach

#### Study design and protocol

A study protocol was created before development of NP-SOLES and includes eligibility criteria for studies, search terms and data to be extracted. Amendments to the protocol were recorded.

#### Systematic searches

To identify relevant neuropathic pain literature, a comprehensive database of studies was assembled from prior human-led systematic searches of PubMed, Web of Science, BIOSIS Citation Index, BIOSIS Previews, and Embase conducted in September 2012 and updated in November 2015. To ensure coverage of more recent literature, additional searches of PubMed, Web of Science, BIOSIS Citation Index, Embase, and Scopus were conducted between November and December 2023. Search strategies were designed to capture animal studies of neuropathic pain and were applied consistently across databases. Retrieved records from all sources were combined into a single corpus, and automated deduplication was used to remove duplicate entries.

#### Machine-Assisted Screening

A machine-assisted screening approach was then applied to support efficient identification of potentially eligible studies from the neuropathic pain corpus. A supervised machine-learning classifier was trained on manually screened studies labelled as eligible (1) or ineligible (0) for inclusion according to predefined eligibility criteria. To update and refine classifier performance, a random sample of 2,000 records retrieved in November 2023 was independently screened at title and abstract level by two reviewers using the SyRF platform (Bahor et al., 2021; Research Resource Identifier (RRID): SCR_018907). Classifier performance was evaluated in terms of sensitivity and specificity for identifying potentially eligible studies, with target thresholds of >0.95 sensitivity and >0.80 specificity.

To further improve classifier performance, we conducted error analysis by selecting 500 studies with the highest level of disagreement between manual and machine screening, and excluded those which lacked an abstract (n=82); i.e. manually screened 418 studies. These studies were re-screened using the same process as before, with discrepancies between reviewers resolved through discussion, and a third reviewer consulted when consensus could not be reached.

### Pilot Systematic Review

#### Study design and protocol

This study comprised a systematic review and meta-analysis of controlled preclinical animal studies evaluating the effects of citrus flavonoids on pain-related outcomes in models of painful diabetic neuropathy (PDN). The review was conducted in accordance with established guidelines for preclinical systematic reviews, with a protocol developed prior to study selection. The protocol was not pre-registered and was not publicly available before the review began. The protocol and full search strategies are provided in the Supplementary Materials.

#### Rationale for outcome selection

Thermal and mechanical pain-related outcomes were selected because both are commonly used behavioural measures in STZ-induced models of painful diabetic neuropathy and reflect distinct nociceptive modalities relevant to clinical symptoms (Sadler, Mogil and Stucky, 2022). Where sufficient data were available, outcomes were synthesised quantitatively by modality, otherwise qualitatively.

#### Study identification

Following application of the NP-SOLES machine learning classifier to the systematic search results, study identification was facilitated through development of regular expressions for text mining within NP-SOLES to automatically tag and filter studies involving citrus flavonoids, streptozotocin-induced animal models of diabetic neuropathic pain, and thermal and mechanical pain outcomes. These codes are provided in Supplementary Materials.

#### Eligibility criteria

Studies were eligible for inclusion if they met the following criteria:

- Population: Non-human animal models of painful diabetic neuropathy induced using streptozotocin (STZ).
- Intervention: Administration of citrus-derived flavonoids, either as isolated compounds or defined extracts.
- Comparator: Vehicle-treated or untreated control groups.
- Outcomes: Quantitative measures of pain-related behaviour, with a primary focus on thermal and mechanical pain-related outcomes (e.g. withdrawal latency or threshold).
- Study design: Controlled experimental studies.

Studies were excluded if they did not report using a model of streptozotocin-induced painful diabetic neuropathy, lacked an appropriate control group, did not report relevant pain-related outcomes, or were not primary research articles.

#### Study selection

Following automated prioritisation, titles and abstracts were screened against the eligibility criteria. Full-text articles were retrieved for studies deemed potentially eligible. Final inclusion was determined based on full-text assessment. The study selection process is summarised using a flow diagram (Figure 1).

**Figure 1.**
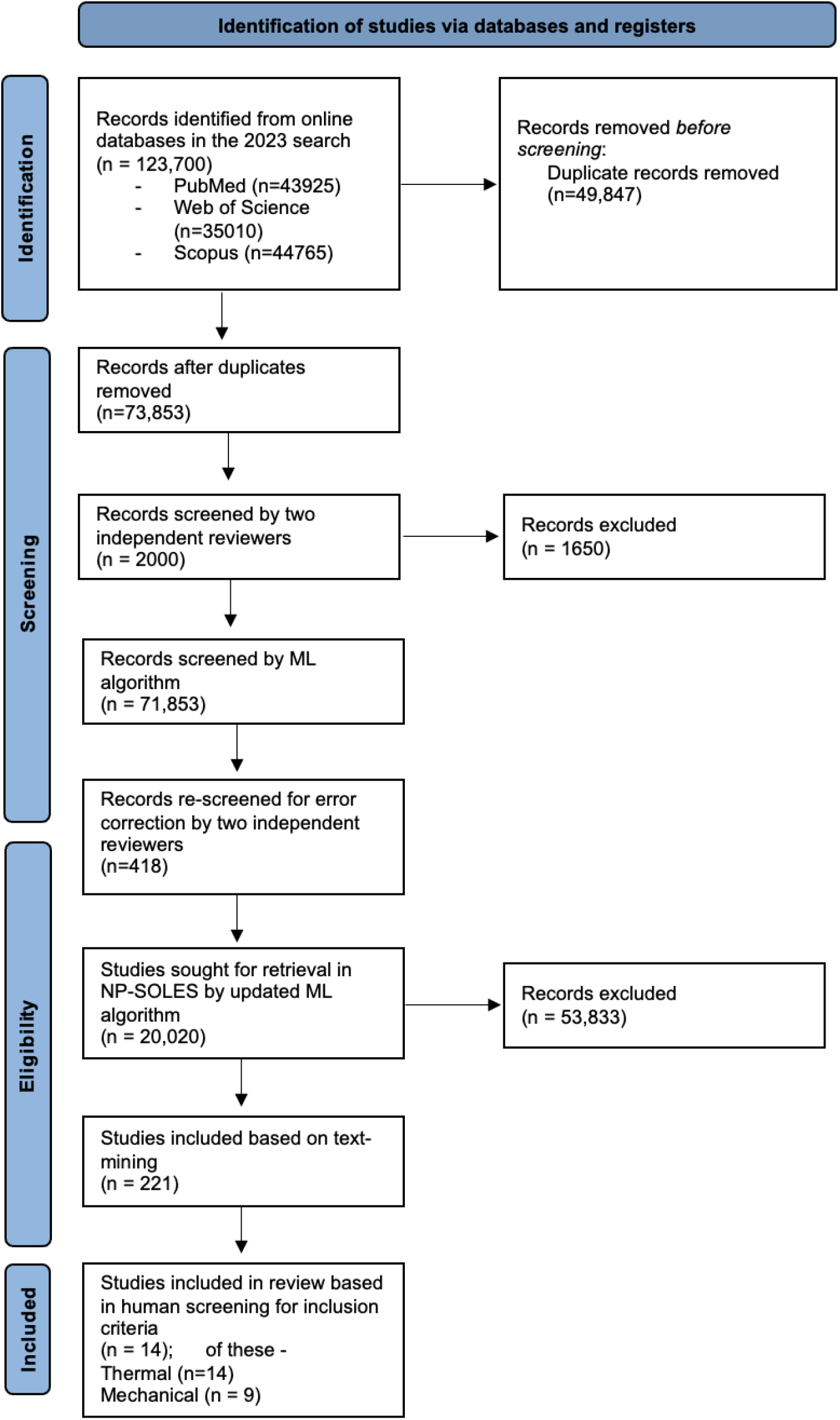
Flow diagram summarising study identification, screening, and inclusion. Initial screening targeted animal studies of neuropathic pain, followed by restriction to streptozotocin-induced painful diabetic neuropathy models assessing citrus flavonoids.

#### Data extraction

Data were extracted from included studies using a predefined extraction form. Extracted information included animal species and strain, sex, induction protocol, intervention characteristics (compound, dose, route, and timing), outcome measures (eg. behavioural test used, site tested, timepoint of data extraction), and numerical data required for meta-analysis (eg. sample size, mean and error values of outcome measurements). Where outcome measurements were taken at several timepoints over a treatment window, the outcome measurement at the longest duration of treatment was taken. Where multiple pain-related outcomes were reported, thermal and mechanical pain-related behavioural outcomes were extracted. Where numerical data were not reported in text or tables, values were extracted from figures, where possible, using WebPlotDigitizer and recorded in SyRF. Following data extraction, all data was exported from SyRF to Microsoft Excel to support data analysis.

#### Risk of bias and reporting quality assessment

Risk of bias and reporting quality were assessed using a modified CAMARADES checklist, including items relating to randomisation, blinding, sample size calculation, and reporting of animal characteristics and husbandry conditions. Each item was recorded as reported or not reported. Risk of bias assessments were used to inform interpretation of findings and exploration of heterogeneity, rather than as exclusion criteria.

#### Data synthesis and statistical analysis

Meta-analysis was conducted where sufficient data were available using a random-effects model to account for between-study heterogeneity. Effect sizes were calculated as standardised mean differences with corresponding 95% confidence intervals. Statistical heterogeneity was assessed using the I^2^ statistic.

Prespecified subgroup analyses explored sources of heterogeneity where data permitted, including outcome type and selected study characteristics. Subgroup analyses were interpreted cautiously, particularly where the number of contributing studies was small. Due to limited reporting of female animals, analyses examining the influence of animal sex were considered exploratory and were not used to draw definitive conclusions.

For the primary analysis, effect sizes from studies contributing multiple flavonoid dose groups against a shared control were nested within study to avoid inflating precision through double-counting of shared control animals. For the subgroup analysis exploring assay type as a source of heterogeneity (Figure 3), this nesting was not applied: because thermal pain assay is a property of the individual outcome measurement rather than the study as a whole, aggregating multiple assays within a study into a single nested effect size would have removed the very distinction this subgroup analysis was designed to examine. This analysis therefore used all individual comparisons at the assay level, and its pooled estimate is consequently not directly comparable to the primary pooled estimate in Figure 2.

**Figure 2.**
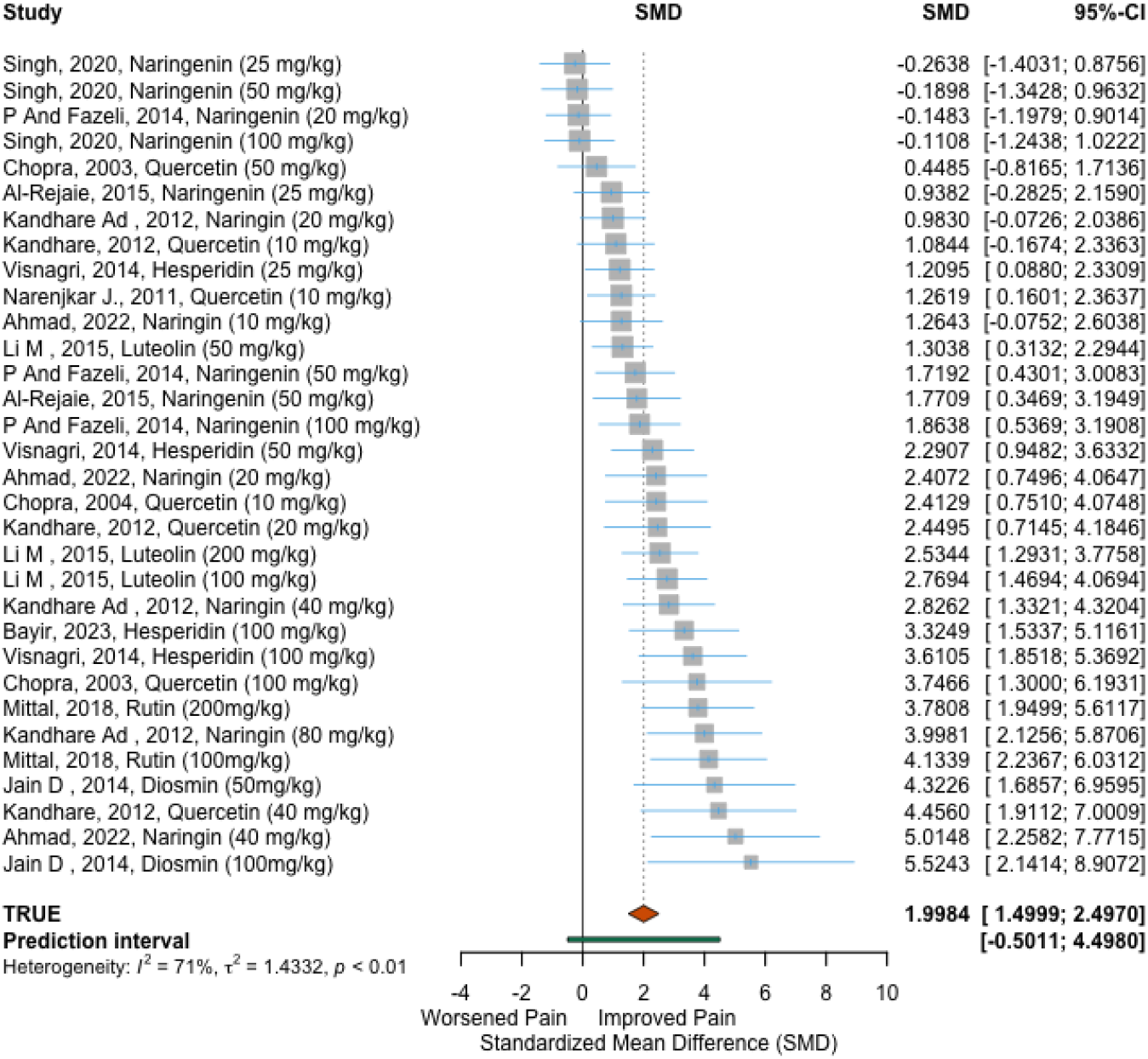
Forest plot showing the effects of citrus flavonoids on thermal pain-related outcomes in streptozotocin-induced models of painful diabetic neuropathy. Effect sizes are presented as standardised mean differences using a random-effects model.

**Figure 3.**
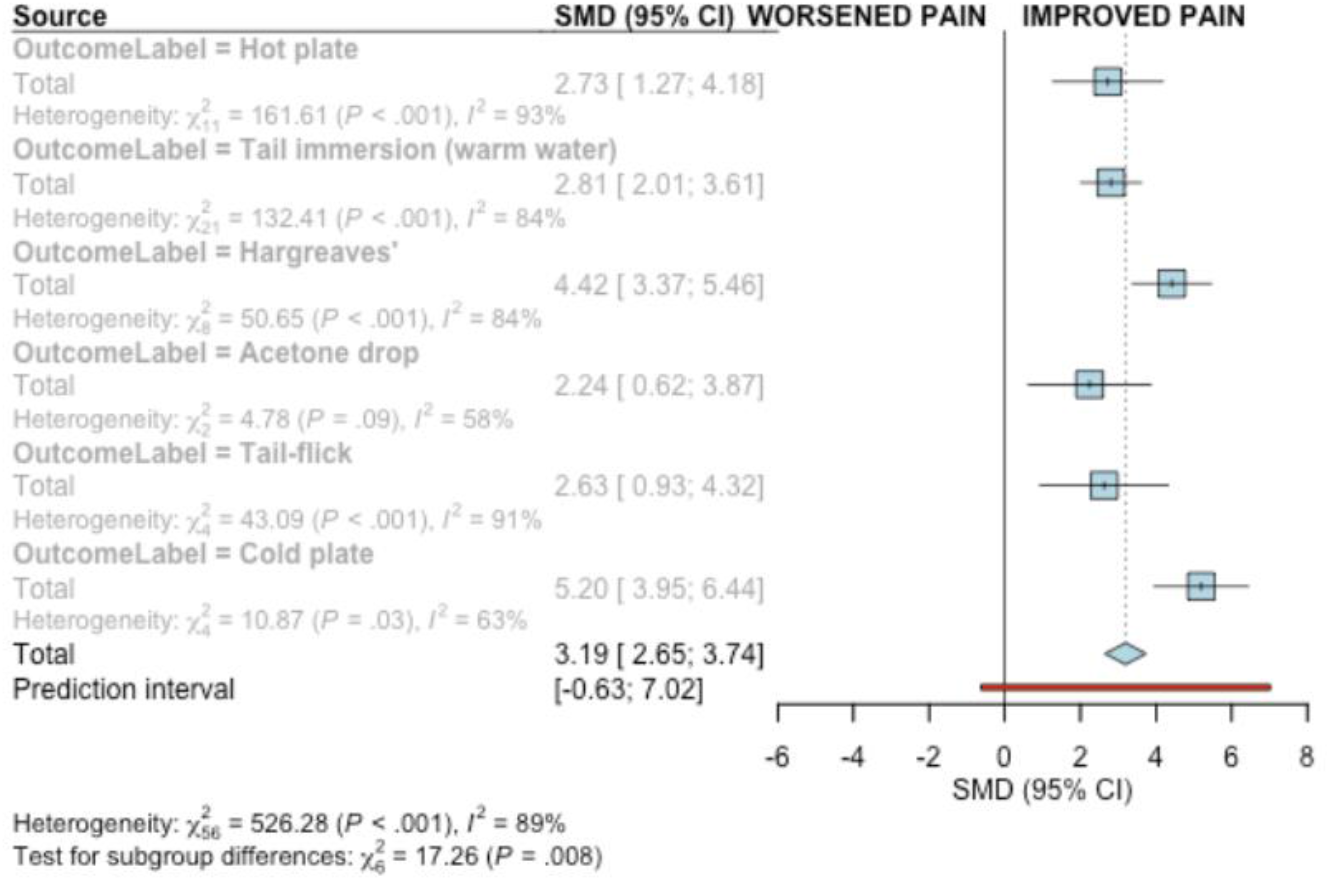
Exploratory subgroup analysis by thermal pain assay. Forest plot showing subgroup analyses of thermal pain-related outcomes stratified by assay type (hot plate, tail immersion, Hargreaves’ test, and related assays). Subgroup estimates are presented for descriptive purposes and should be interpreted cautiously due to limited numbers of contributing studies and substantial heterogeneity.

All statistical analyses were conducted using R (version 4.3.3; R Core Team, 2024). Data were imported and processed using readxl (v1.4.3) and the tidyverse (v2.0.0; Wickham et al., 2019). Standardised mean differences were calculated following Vesterinen et al. (2014), and random-effects meta-analysis (REML estimator of τ^2^), subgroup analyses and meta-regression were performed using meta (v7.0.0; Balduzzi et al., 2019) and metafor (v4.6.0; Viechtbauer, 2010). Additional methodological details and analysis scripts are provided in the Supplementary Materials.

#### Role of automation in analysis

Machine-assisted methods were used solely to support study identification and screening. Data extraction, risk of bias assessment, and statistical analysis were conducted manually.

## Results

### Refinement of the machine learning-classifier

Error analysis and refinement of the machine-learning classifier showed an improvement in the performance of the algorithm with specificity changing from 63% to 73% and sensitivity changing from 96% to 95%. Despite the improvement in specificity, we were still unable to meet the 80% target that we were aiming for.

### Study selection

Application of the machine-assisted screening approach enabled identification of 221 potentially relevant studies from the large neuropathic pain literature database (NP-SOLES). Following automated prioritisation and manual screening, full texts were assessed for eligibility. Reasons for exclusion at the full-text stage included not using a model of streptozotocin-induced painful diabetic neuropathy-, absence of citrus flavonoid interventions, lack of appropriate control groups, or failure to report relevant pain-related outcomes. The study selection process is summarised in Figure 1. Fourteen studies meeting the eligibility criteria reported thermal pain related outcomes and nine studies of the 14 reported mechanical pain-related outcomes.

#### Characteristics of included studies

The fourteen included studies used streptozotocin (STZ)-induced models of diabetic neuropathy across a range of animal species and strains (Table 1). Citrus flavonoids were administered using varied compounds, doses, routes, and treatment schedules. Studies assessed pain-related behaviour using thermal assays, including measures of withdrawal latency or threshold.

**Table 1.** Study characteristics of the included studies.

| Author (Year) and DOI | Species | Sex | STZ dose (mg/kg) | Citrus flavonoid | Dose (s) (mg/kg) | Duration (weeks) | Test(s) |
| --- | --- | --- | --- | --- | --- | --- | --- |
| Ahmad *et al.* (2022)<br>doi:https://doi.org/10.3390/life12122111 | Wistar rats | Male | 35 | Naringin | 10; 20; 40 | 4 | Acetone drop; Hot plate; Tail immersion (warm); Plantar test |
| Hasanein & Fazeli (2014)<br>doi:https://doi.org/10.1007/s13105-014-0369-5 | Wistar rats | Male | 50 | Naringenin | 20; 50; 100 | 8 | Tail flick; Von Frey |
| Mittal *et al.* (2018)<br>doi:https://doi.org/10.1007/s10787-017-0413-5 | Wistar rats | Male | 55 | Rutin | 100; 200 | 8 | Cold plate; Hot plate; Von Frey & Randall-Selitto |
| Bayir *et al.* (2023)<br>doi:https://doi.org/10.1007/s12640-023-00657-0 | Wistar rats | Not reported | 45 | Hesperidin | 100 | 2 | Hot plate; Tail immersion (warm); |
| Anjaneyulu & Chopra (2003)<br>doi:[https://doi.org/10.1016/S0278-5846(03)00160-X](https://doi.org/10.1016/S0278-5846%2803%2900160-X) | Laka mice | Male | 200 | Quercetin | 50; 100 | N/A | Tail immersion (warm) |
| Anjaneyulu & Chopra (2004) | Sprague Dawley rats | Male | 45 | Quercetin | 10 | 4 | Tail immersion (warm); Tail immersion (cold); Hot plate |
| Narenjkar *et al.* (2011) | Wistar rats | Male | 60 | Quercetin | 10 | 6 | Tail immersion (warm) |
| Kandhare *et al.* (2012a)<br>doi:https://doi.org/10.1016/j.fitote.2012.01.010 | Wistar rats | Male | 55 | Naringin | 20; 40; 80 | 4 | Tail immersion (warm); Von Frey & Randall-Selitto |
| Jain *et al.* (2014)<br>doi:[https://doi.org/10.1016/s2095-4964(14)60001-7](https://doi.org/10.1016/s2095-4964%2814%2960001-7) | Sprague Dawley rats | Male | 35 | Diosmin | 50; 100 | 4 | Hot plate; Tail immersion (cold) |
| Kandhare *et al.* (2012b)<br>doi:https://doi.org/10.1016/j.biomag.2012.10.002 | Wistar rats | Both | 90 | Quercetin | 10; 20; 40 | 8 | Hargreaves' test; Tail immersion (warm); Von Frey & Randall-Selitto |
| Al-Rejaie *et al.* (2015)<br>doi:https://doi.org/10.1179/1743132815y.0000000079 | Wistar rats | Male | 60 | Naringenin | 25; 50 | 5 | Tail flick; Randall-Selitto |
| Visnagri *et al.* (2014)<br>doi:https://doi.org/10.3109/13880209.2013.870584 | Sprague<br>Dawley rats | *Not reported* | 55 | Hesperidin | 25;<br>50;<br>100 | 4 | Hargreaves' test;<br>Tail immersion (warm);<br>Von Frey & Randall-Selitto |
| Singh *et al.* (2020)<br>doi:https://doi.org/10.1039/c9fo00881k | Wistar rats | *Not reported* | 45 | Naringenin | 25;<br>50;<br>100 | 4 | Tail immersion (warm);<br>Hot plate;<br>Von Frey & Randall-Selitto |
| Li *et al.* (2015) | Sprague<br>Dawley rats | Male | 55 | Luteolin | 50;<br>100;<br>200 | 3 | Hargreaves' test;<br>Cold plate;<br>Von Frey |

Reporting of animal sex was inconsistent. Most studies used male animals, with only one study reporting outcomes in females. As a result, sex-specific analyses were limited.

#### Thermal Pain-Related Outcomes

Thermal pain-related outcomes were analysed using quantitative methods and mechanical pain-related outcomes were analysed using qualitative methods according to pre-defined criteria.

### Effects of citrus flavonoids on thermal pain-related outcomes

Meta-analysis of thermal pain-related outcomes suggested a reduction in pain-related behaviours following treatment with citrus flavonoids compared with control conditions (Figure 2). Effect size estimates varied substantially across studies, indicating marked between-study heterogeneity.

While the pooled estimate indicated a potential analgesic effect, individual study results ranged from modest to large effects, and confidence intervals were wide in several cases, reflecting small sample sizes and variability in experimental design.

### Heterogeneity and subgroup analyses

Statistical heterogeneity was substantial (I^2^ indicating considerable between-study variability). Prespecified subgroup analyses explored potential sources of heterogeneity where data permitted, including differences in outcome assessment and selected study characteristics.

Due to limited reporting and small numbers of contributing studies, subgroup analyses were underpowered and were interpreted cautiously. Although animal sex was explored as a potential source of heterogeneity, meaningful conclusions could not be drawn given that only a single study reported female data. As such, any apparent differences by sex should be considered exploratory.

#### Mechanical Pain-Related Outcomes

Nine studies reported mechanical pain-related outcomes, most commonly tactile allodynia assessed using von Frey-based methods. Across studies, treatment with citrus flavonoids was associated with increased mechanical withdrawal thresholds or reduced mechanical hypersensitivity at one or more tested doses compared with control conditions.

However, substantial heterogeneity was observed in animal characteristics, neuropathy induction protocols, outcome measures, and dosing regimens. Mechanical outcomes were not consistently reported across studies, and differences in testing paradigms and data presentation precluded quantitative synthesis. As less than 10 studies reported mechanical outcomes, a meta-analysis was not conducted in keeping with the protocol. As such, findings for mechanical pain-related outcomes are based on qualitative analysis only. Detailed study-level results are provided in the Supplementary Materials.

#### Risk of bias and reporting quality

Reporting of key measures intended to reduce risk of bias such as randomisation, blinding, and sample size calculation were poor (Fig 4), highlighting widespread methodological limitations. Animal welfare compliance such as seeking ethical approval was successfully carried out across all studies. Details regarding animal housing, husbandry, and experimental conditions were frequently omitted.

**Figure 4.**
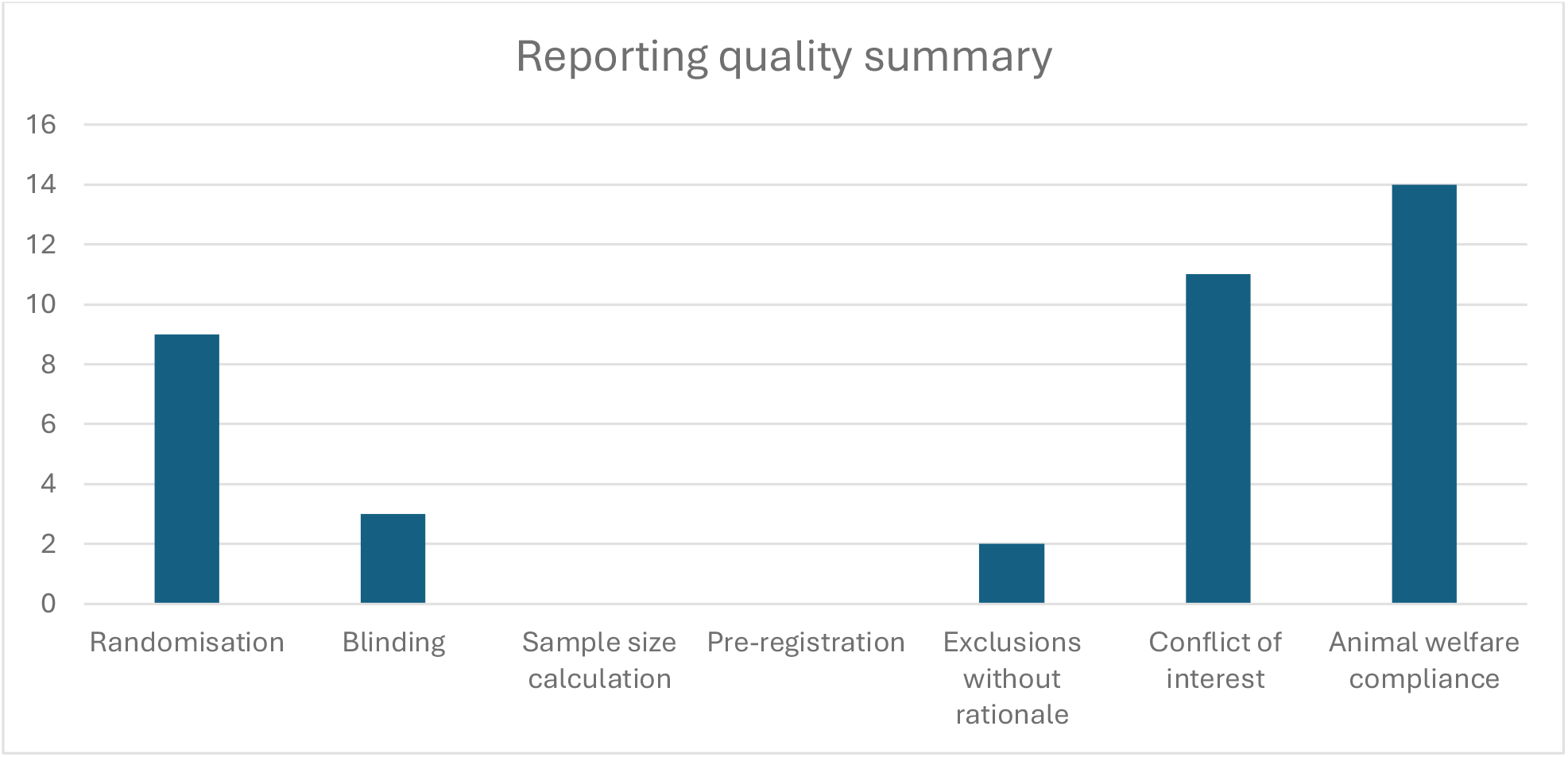
Reporting quality and risk of bias indicators across included studies. Bars show the number of included studies reporting key methodological and transparency items based on a modified CAMARADES checklist

The limitations identified were consistent across studies and were considered likely contributors to the observed heterogeneity and to potential inflation of treatment effect estimates.

## Discussion

In this study, we evaluated the effects of citrus flavonoids on thermal and mechanical pain-related outcomes in streptozotocin-induced models of painful diabetic neuropathy using a machine-assisted screening approach. Pooled analyses suggested a reduction in thermal pain-related behaviours and qualitative analysis indicates potential reduction in mechanical pain-related behaviours; however, substantial heterogeneity and widespread limitations in study design and reporting as well a lack of controlled experimental data on mechanical pain-related outcomes constrain confidence in these findings. Together, these results highlight both the potential promise of citrus flavonoids as candidate interventions and the persistent challenges facing preclinical neuropathic pain research.

### Interpretation of findings

#### Thermal pain-related outcomes

The observed reduction in thermal pain-related outcomes across studies is consistent with reports from individual animal experiments suggesting analgesic effects of citrus flavonoids. Proposed biological mechanisms include antioxidant and anti-inflammatory actions, which are plausible given the established roles of oxidative stress and neuroinflammation in the pathophysiology of diabetic neuropathy. However, the magnitude and consistency of the observed effects varied considerably between studies, and pooled estimates were accompanied by wide confidence and prediction intervals.

Subgroup analyses stratified by thermal pain assay indicated variability in effect estimates across outcome measures, suggesting that differences in behavioural testing paradigms may contribute to heterogeneity. Heterogeneity in experimental design and outcome assessment is not inherently problematic and may, in some contexts, strengthen confidence in the generalisability of preclinical findings if treatment effects are observed consistently across models and assays. However, in the present literature, variability in outcome assessment co-occurred with poor reporting of key methodological safeguards, making it difficult to distinguish meaningful biological differences from nuisance variability introduced by experimental design, analysis choices, or selective reporting. Under such conditions, heterogeneity may obscure true effects or contribute to inflated estimates of treatment efficacy (Voelkl and Würbel,2024). It therefore remains unclear whether differences in observed effects across assays reflect distinct underlying nociceptive mechanisms or arise primarily from uncontrolled methodological factors.

Subgroup analyses were exploratory and underpowered and should not be interpreted as evidence of differential efficacy across assays.

#### Mechanical pain-related outcomes

Across included studies, citrus flavonoid treatment was generally associated with reductions in mechanical pain-related behaviours compared with control conditions. However, interpretation of these findings is constrained by substantial variability in experimental design and outcome measurement. Differences in animal models of diabetic neuropathy resulted in heterogeneous baseline sensitivities, while factors such as housing conditions-that may have roles in influencing stress and behavioural responses (Arndt *et al*., 2009) were inconsistently reported.

Citrus flavonoid interventions also varied markedly with respect to compound, dose, treatment duration, and dosing frequency. Although some studies reported analgesic effects only at the upper end of their own dosing ranges, often described in studies as “higher doses”, no consistent dose threshold could be identified to classify a dose as high or low, and definitions of “low” and “high” doses were not provided but rather described as relative to doses used in the experiments. Given that less than 10 studies reported mechanical pain-related outcomes, evidence for these is based on qualitative analysis only and should therefore be interpreted cautiously.

### Risk of bias, reporting quality, and implications for translation

A key finding of this review was the poor reporting of measures intended to reduce risk of bias across included studies. Blinding, sample size calculation, and preregistration were infrequently reported, consistent with previous assessments of preclinical pain literature. Randomisation although more frequently reported, the methods were not disclosed raising the potential doubts on its validity. Such limitations are known to be associated with inflated effect size estimates and reduced reproducibility, raising the possibility that the apparent analgesic effects observed here may overestimate true efficacy. Compliance with animal welfare regulations was widely reported and all studies reported obtaining ethical approval.

Sex-specific analyses were severely limited by the predominance of male-only studies and inconsistent reporting of animal sex. A similar trend can be seen across neuroscience research. Given increasing recognition of sex differences in pain mechanisms and treatment response, this represents a significant gap in the evidence base and further limits translational relevance posing detrimental effects to patient populations especially in female predominant conditions such as pain disorders (Hurley and Adams, 2008).

The method of PDN modelling using streptozotocin also needs to be considered when thinking about translation to clinical research. Although it is currently the most common choice of diabetes inducing agent for PDN, its toxic effects on multiple organ systems at the high doses used for diabetes induction and the lack of representation of the chronic progressive and heterogenic nature of diabetes in humans must be considered (Furman, 2021).

Taken together, these findings underscore the need for more rigorously designed and transparently reported preclinical studies before citrus flavonoids can be considered credible candidates for clinical investigation in painful diabetic neuropathy.

### Role of machine-assisted evidence synthesis

This review also demonstrates the feasibility of applying machine-assisted screening approaches to support preclinical systematic reviews in neuropathic pain research. The use of automated deduplication, text-based filtering, and machine learning-assisted prioritisation reduced manual screening burden while maintaining human oversight of all inclusion decisions. Although this study represents a pilot application rather than a comprehensive evaluation of the screening approach, it illustrates how such methods may help address scalability challenges in evidence synthesis as the literature continues to expand.

Further development and validation of machine-assisted tools across a broader range of neuropathic pain models and outcomes is ongoing. Importantly, automation should be viewed as a complement to, rather than a replacement for, rigorous human judgement in systematic review methodology.

### Limitations

This review has several limitations. First, the number of included studies was modest and heterogeneity across experimental designs was substantial, limiting the precision of pooled effect estimates. For mechanical pain-related outcomes in particular, the number and consistency of reported measures were insufficient to support quantitative synthesis, and findings were therefore restricted to descriptive analysis.

Second, our approach to handling non-independent comparisons was not fully consistent across analyses. In the primary meta-analysis, effect sizes from studies contributing multiple dose groups against a shared control were nested within study to avoid double-counting shared control animals. The subgroup analysis exploring pain assay type as a source of heterogeneity (Figure 3), however, required disaggregating these nested comparisons, since assay type is a property of the individual outcome measurement rather than the study as a whole; this analysis therefore used non-nested, individual comparison-level data, and its pooled estimate is not directly comparable to the primary analysis. A multi-level meta-regression model capable of accounting for both shared controls and assay-level heterogeneity simultaneously would have offered a more statistically consistent treatment of these data but was beyond the scope of this pilot review.

Third, although machine-assisted screening substantially reduced manual workload, it did not identify all relevant studies when compared with supplementary manual searching, raising the possibility that some eligible evidence was missed. In addition, the definition of “citrus flavonoids” required refinement during protocol development due to heterogeneity in how compounds were described in the literature, which may have influenced study inclusion.

Fourth, restriction of outcomes to thermal and mechanical pain-related behaviours, while methodologically justified, excluded other clinically relevant modalities and more complex behavioural assessments that may better capture aspects of animal pain and wellbeing. Analyses were further constrained by incomplete reporting of key study characteristics, including animal sex and husbandry conditions, limiting exploration of potential sources of variability.

Finally, while this work demonstrates the feasibility and potential value of machine-assisted systematic review approaches in preclinical neuropathic pain research, it was not designed to formally evaluate the performance of automated screening relative to fully manual methods. As such, this study should be considered a proof-of-concept illustrating the application of NP-SOLES rather than a definitive assessment of the platform’s capabilities.

## Conclusions

Available preclinical evidence suggests that citrus flavonoids may reduce thermal and mechanical pain-related behaviours, in streptozotocin-induced models of painful diabetic neuropathy. However, confidence in these findings is limited by substantial heterogeneity, limited high-quality evidence, and pervasive risks of bias across the underlying studies. Improved methodological rigor and transparency in reporting are required to clarify the robustness and translational relevance of these effects. More broadly, this work demonstrates the potential of machine-assisted approaches to support timely and scalable synthesis of preclinical evidence in neuropathic pain research.

## Supporting information

SyRCLE protocol

Mechanical outcomes summary

R script for Meta Analysis

Regular expression codes

Thermal outcomes summary

## Author Contributions (CRediT)

S.J.: Conceptualization, Formal analysis, Investigation, Data curation, Writing - original draft, Writing - review & editing.

J.M.: Conceptualization, Investigation, Data curation, Writing – original draft, Writing - review & editing.

N.S.: Conceptualization, Writing-reviewing & editing.

S.S.: Software, Methodology.

G.C.: Conceptualization, Methodology, Supervision, Writing - review & editing.

## Supplementary Materials

Protocol for Pilot Systematic Review

Regular Expression codes

R script for meta analysis

Pilot Systematic Review Detailed Findings

## References

Arndt, S. S., Laarakker, M. C., van Lith, H. A., van der Staay, F. J., Gieling, E., Salomons, A. R., van’t Klooster, J. and Ohl, F. (2009) ‘Individual housing of mice—impact on behaviour and stress responses’, Physiology & behavior, 97(3-4), pp. 385–393.

Cavalli, E., Mammana, S., Nicoletti, F., Bramanti, P. and Mazzon, E. (2019) ‘The neuropathic pain: An overview of the current treatment and future therapeutic approaches’, International journal of immunopathology and pharmacology, 33, pp. 2058738419838383.

Colloca, L., Ludman, T., Bouhassira, D., Baron, R., Dickenson, A. H., Yarnitsky, D., Freeman, R., Truini, A., Attal, N. and Finnerup, N. B. (2017) ‘Neuropathic pain’, Nature reviews Disease primers, 3(1), pp. 1–19.

Finnerup, N. B., Kuner, R. and Jensen, T. S. (2020) ‘Neuropathic pain: from mechanisms to treatment’, Physiological reviews.

Fisher, A. S., Lanigan, M. T., Upton, N. and Lione, L. A. (2021) ‘Preclinical neuropathic pain assessment; the importance of translatability and bidirectional research’, Frontiers in Pharmacology, 11, pp. 614990.

Fox, A., Eastwood, C., Gentry, C., Manning, D. and Urban, L. (1999) ‘Critical evaluation of the streptozotocin model of painful diabetic neuropathy in the rat’, Pain, 81(3), pp. 307–316.

Furman, B. L. (2021) ‘Streptozotocin‐induced diabetic models in mice and rats’, Current protocols, 1(4), pp. e78.

Hurley, R. W. and Adams, M. C. (2008) ‘Sex, gender, and pain: an overview of a complex field’, Anesthesia & Analgesia, 107(1), pp. 309–317.

Ineichen, B. V., Held, U., Salanti, G., Macleod, M. R. and Wever, K. E. (2024) ‘Systematic review and meta-analysis of preclinical studies’, Nature Reviews Methods Primers, 4(1), pp. 72.

Jensen, T. S., Baron, R., Haanpää, M., Kalso, E., Loeser, J. D., Rice, A. S. and Treede, R.-D. (2011) ‘A new definition of neuropathic pain’, Pain, 152(10), pp. 2204–2205.

Jin, H. Y., Moon, S.-S. and Calcutt, N. A. (2021) ‘Lost in translation? Measuring diabetic neuropathy in humans and animals’, Diabetes & metabolism journal, 45(1), pp. 27–42.

Mulrow, C. D. (1994) ‘Systematic reviews: rationale for systematic reviews’, Bmj,309(6954), pp. 597–599.

Sadler, K. E., Mogil, J. S. and Stucky, C. L. (2022) ‘Innovations and advances in modelling and measuring pain in animals’, Nature Reviews Neuroscience, 23(2), pp. 70–85.

van Velzen, M., Dahan, A. and Niesters, M. (2020) ‘Neuropathic pain: challenges and opportunities’, Frontiers in Pain Research, 1, pp. 1.

Vincent, A. M., Callaghan, B. C., Smith, A. L. and Feldman, E. L. (2011) ‘Diabetic neuropathy: cellular mechanisms as therapeutic targets’, Nature Reviews Neurology, 7(10), pp. 573–583.

Voelkl, B. and Würbel, H. (2024) ‘Heterogeneity of animal experiments and how to deal with it’, Laboratory Animals, 58(5), pp. 493–497.

Wang, Y., Liu, X.-J., Chen, J.-B., Cao, J.-P., Li, X. and Sun, C.-D. (2022) ‘Citrus flavonoids and their antioxidant evaluation’, Critical Reviews in Food Science and Nutrition, 62(14), pp. 3833–3854.

## References of Included Studies

Ahmad, M. F., Naseem, N., Rahman, I., Imam, N., Younus, H., Pandey, S. K. and Siddiqui, W. A. (2022) ‘Naringin attenuates the diabetic neuropathy in STZ-induced type 2 diabetic Wistar rats’, Life, 12(12), p. 2111. doi: 10.3390/life12122111.

Al-Rejaie, S. S., Aleisa, A. M., Abuohashish, H. M., Parmar, M., Ola, M. S., Al-Hosaini, A.A. and Ahmed, M. M. (2015) ‘Naringenin neutralises oxidative stress and nerve growth factor discrepancy in experimental diabetic neuropathy’, Neurological Research, 37(10), pp. 924–933. doi: 10.1179/1743132815Y.0000000079.

Anjaneyulu, M. and Chopra, K. (2003) ‘Quercetin, a bioflavonoid, attenuates thermal hyperalgesia in a mouse model of diabetic neuropathic pain’, Progress in Neuro-Psychopharmacology & Biological Psychiatry, 27(6), pp. 1001–1005. doi: 10.1016/S0278-5846(03)00160-X.

Anjaneyulu, M. and Chopra, K. (2004) ‘Quercetin attenuates thermal hyperalgesia and cold allodynia in STZ-induced diabetic rats’, Indian journal of experimental biology, 42(8), pp. 766–769.

Balduzzi S, Rücker G, Schwarzer G (2019), How to perform a meta-analysis with R: a practical tutorial, Evidence-Based Mental Health; 22:153–160.

Bayir, M. H., Yıldızhan, K. and Altındağ, F. (2023) ‘Effect of hesperidin on sciatic nerve damage in STZ-induced diabetic neuropathy: modulation of TRPM2 channel’, Neurotoxicity Research, 41(6), pp. 638–647. doi: 10.1007/s12640-023-00657-0.

Hasanein, P. and Fazeli, F. (2014) ‘Role of naringenin in protection against diabetic hyperalgesia and tactile allodynia in male Wistar rats’, Journal of Physiology and Biochemistry, 70(4), pp. 997–1006. doi: 10.1007/s13105-014-0369-5.

Jain, D., Bansal, M. K., Dalvi, R., Upganlawar, A. and Somani, R. (2014) ‘Protective effect of diosmin against diabetic neuropathy in experimental rats’, Journal of Integrative Medicine, 12(1), pp. 35–41. doi: 10.1016/S2095-4964(14)60001-7.

Kandhare, A. D., Raygude, K. S., Ghosh, P., Ghule, A. E. and Bodhankar, S. L. (2012a) ‘Neuroprotective effect of naringin by modulation of endogenous biomarkers in streptozotocin-induced painful diabetic neuropathy’, Fitoterapia, 83(4), pp. 650–659. doi: 10.1016/j.fitote.2012.01.010.

Kandhare, A. D., Raygude, K. S., Kumar, V., Rajmane, A. R., Visnagri, A., Ghule, A. E., Ghosh, P., Badole, S. L. and Bodhankar, S. L. (2012b) ‘Ameliorative effects of quercetin against impaired motor nerve function, inflammatory mediators and apoptosis in neonatal streptozotocin-induced diabetic neuropathy in rats’, Biomedicine & Aging Pathology, 2(4), pp. 173–186. doi: 10.1016/j.biomag.2012.10.002.

Li, M., Li, Q., Zhao, Q., Zhang, J. and Lin, J. (2015) ‘Luteolin improves the impaired nerve functions in diabetic neuropathy: behavioral and biochemical evidences’, International Journal of Clinical and Experimental Pathology, 8(9), pp. 10112–10120.

Mittal, R., Kumar, A., Singh, D. P., Bishnoi, M. and Nag, T. C. (2018) ‘Ameliorative potential of rutin in combination with nimesulide in STZ model of diabetic neuropathy: targeting Nrf2/HO-1/NF-kB and COX signalling pathway’, Inflammopharmacology, 26(3), pp. 755–768. doi: 10.1007/s10787-017-0413-5.

Narenjkar, J., Roghani, M., Alambeygi, H. and Sedaghati, F. (2011) ‘The effect of the flavonoid quercetin on pain sensation in diabetic rats’, Basic and Clinical Neuroscience, 2(3), pp. 51–57.

R Core Team (2024). _R: A Language and Environment for Statistical Computing_. R Foundation for Statistical Computing, Vienna, Austria. <https://www.R-project.org/>.

Singh, P., Bansal, S., Kuhad, A., Kumar, A. and Chopra, K. (2020) ‘Naringenin ameliorates diabetic neuropathic pain by modulation of oxidative-nitrosative stress, cytokines and MMP-9 levels’, Food & Function, 11(5), pp. 4548–4560. doi: 10.1039/C9FO00881K.

Viechtbauer, W. (2010). Conducting meta-analyses in R with the metafor package. Journal of Statistical Software, 36(3), 1–48. 10.18637/jss.v036.i03

Visnagri, A., Kandhare, A. D., Chakravarty, S., Ghosh, P. and Bodhankar, S. L. (2014) ‘Hesperidin, a flavanoglycone attenuates experimental diabetic neuropathy via modulation of cellular and biochemical marker to improve nerve functions’, Pharmaceutical Biology, 52(7), pp. 814–828. doi: 10.3109/13880209.2013.870584.

Wickham H, Averick M, Bryan J, Chang W, McGowan LD, François R, Grolemund G,Hayes A, Henry L, Hester J, Kuhn M, Pedersen TL, Miller E, Bache SM, Müller K, Ooms J,Robinson D, Seidel DP, Spinu V, Takahashi K, Vaughan D, Wilke C, Woo K, Yutani H (2019). “Welcome to the tidyverse.” Journal of Open Source Software, 4(43), 1686. doi:10.21105/joss.01686 https://doi.org/10.21105/joss.01686.

