## Supplementary material for "Machine-assisted systematic review and meta-analysis of citrus flavonoids in streptozotocin-induced painful diabetic neuropathy": SyRCLE protocol

### SYSTEMATIC REVIEW PROTOCOL FOR ANIMAL INTERVENTION STUDIES

FORMAT BY SYRCLE (WWW.SYRCLE.NL)
VERSION 2.0 (DECEMBER 2014)

| Item # | Section/Subsection/Item | Description | Check for approval |
|---|---|---|---|
| | A. General | | |
| 1. | Title of the review | Efficacy of Citrus Flavonoids in Alleviating Thermal and Mechanical Pain Behavioural Outcomes: A Systematic Review in the STZ-Induced Diabetic Neuropathy Model | |
| 2. | Authors (names, affiliations, contributions) | Shreya Jindal, Jayashrii Muthaiyan | |
| 3. | Other contributors (names, affiliations, contributions) | Dr. Gillian Currie | |
| 4. | Contact person + e-mail address | | |
| 5. | Funding sources/sponsors | CAMARADES | |
| 6. | Conflicts of interest | None | |
| 7. | Date and location of protocol registration | | |
| 8. | Registration number (if applicable) | | |
| 9. | Stage of review at time of registration | | |
| | B. Objectives | | |
| | Background | | |
| 10. | What is already known about this disease/model/intervention? Why is it important to do this review? | s<br><br>Many animal models have been established to study painful diabetic peripheral neuropathy. The most common is the streptozotocin (STZ)-induced model in rats and mice (Courteix, Eschalier and Lavarenne, 1993). STZ, a glucosamine-nitrosourea compound, is selectively toxic to the insulin-producing beta cells in the pancreas (Junod et al, 1967). This allows the STZ model to replicate key aspects of diabetes, such as hyperglycaemia and insulin deficiency (Ghasemi and Jeddi, 2023). Furthermore, the progression of neuropathic complications in STZ models is similar to human diabetic neuropathy, validating its use in examining the underlying mechanisms of the disease and testing potential therapeutics – particularly in terms of pain-related behaviours.<br><br>Current therapy for DPN focuses on improving glycemic control and analgesia. However, it is far from effective and only delays the onset/progression of disease at best. Preclinical research has highlighted the potential | |

| | | |
|---|---|---|
| | | therapeutic effect of many compounds based on probably pathogenesis, which has led to several clinical trials, but many have failed (Feldman et al, 2019). This limitation is partly due to incomplete understanding of the complexities causing PDN. Therefore, there remains an unmet need for additional therapies that target the underlying causes of the condition.<br><br>There is an increasing interest in exploring natural compounds as alternatives or complements to traditional pharmacological treatments for neuropathic pain. Citrus flavonoids are naturally occurring compounds found in various plants and dietary sources, including citrus fruits. They have garnered attention in the scientific community for their potential therapeutic effects, particularly in the context of neuropathic pain. The interest in citrus flavonoids stems from their wide range of biological properties.<br><br>Citrus flavonoids have been found to reduce oxidative stress, improve glucose tolerance and insulin sensitivity, regulate lipid metabolism and adipocyte differentiation, suppress inflammation and apoptosis, and improve endothelial dysfunction (Mahmoud et al, 2019; Li and Schluesener, 2017; Millar, Duclos and Blesso, 2017; Rees, Dodd and Spencer, 2018; Zaidun, Thent and Abd Latiff, 2018; Zhang et al, 2019; Kopustinskiene et al, 2020). These properties suggest that they could be effective in treating diabetes and potentially reducing the pain associated with diabetic neuropathy.<br><br>While individual studies have explored the effects of citrus flavonoids on diabetic neuropathy, there is a noticeable gap in the systematic consolidation and synthesis of this evidence. This makes it challenging for healthcare professionals and researchers to draw comprehensive conclusions about their efficacy.<br><br>This review aims to fill this gap by systematically compiling, analysing, and summarizing existing research and identify any gaps in the understanding of their effects on different pain behaviours in diabetic neuropathy. This would provide a clearer picture of the efficacy of citrus flavonoids in diabetic neuropathy, potentially guiding future research and clinical practice. |
| | Research question | |
| 11. | Specify the disease/health problem of interest | Painful Diabetic Neuropathy |
| 12. | Specify the population/species studied | STZ-induced animal models |
| 13. | Specify the intervention/exposure | Citrus flavonoids *(Specify)* |

| | | |
|---|---|---|
| 14. | Specify the control population | |
| 15. | Specify the outcome measures | Reflexive Pain tests<br>- Thermal Pain Response - latency or threshold to withdrawal from a thermal stimulus (e.g., hot plate test, tail-flick test). |
| 16. | State your research question (based on items 11-15) | What is the effect of citrus flavonoids in alleviating temperature-related pain-like behaviour in STZ-induced diabetic neuropathy? |
| | C. Methods | |
| | Search and study identification | |
| 17. | Identify literature databases to search (*e.g.* Pubmed, Embase, Web of science) | ☐MEDLINE via PubMed ☐Web of Science<br>☐SCOPUS ☐EMBASE<br>☒Other, namely: NP-SOLES database<br>☐Specific journal(s), namely: |
| 18. | Define electronic search strategies (*e.g.* use the step by step search guide[15] and animal search filters[20, 21]) | When available, please add a supplementary file containing your search strategy: [insert file name] |
| 19. | Identify other sources for study identification | ☐Reference lists of included studies ☐Books<br>☐Reference lists of relevant reviews<br>☐Conference proceedings, namely:<br>☐Contacting authors/ organisations, namely:<br>☐Other, namely: |
| 20. | Define search strategy for these other sources | |
| | Study selection | |
| 21. | Define screening phases (*e.g.* pre-screening based on title/abstract, full text screening, both) | Pre-screening based on title/abstract followed by full text screening for data extraction |
| 22. | Specify (a) the number of reviewers per screening phase and (b) how discrepancies will be resolved | 2 reviewers per screening phase<br>Discrepancies resolved with a third expert party (Dr. Gillian Currie) |
| | *Define all inclusion and exclusion criteria based on:* | |
| 23. | Type of study (design) | Inclusion criteria:<br>All primary experiments in STZ models with appropriate (vehicle/saline) controls<br><br>Exclusion criteria:<br>All studies without appropriate control cohorts and non-primary studies including commentaries, reviews, letters, and editorials |
| 24. | Type of animals/population (*e.g.* age, gender, disease model) | Inclusion criteria:<br>Animal models of streptozotocin induced diabetic neuropathy<br><br>Exclusion criteria: Primary experiments in non-STZ models; studies in humans, *in vitro, ex vivo, or in silico* |

| | | |
|---|---|---|
| 25. | Type of intervention (*e.g.* dosage, timing, frequency) | Inclusion criteria:<br>Citrus flavonoids-<br>- 3,3’,4’,5,6,7,8-Heptamethoxyflavone<br>- 3’,4’,5,7,8-Pentamethoxyflavone<br>- 3,3’,4’,5,7,8-Hexamethoxyflavone<br>- 4’,5,7,8-Tetramethoxyflavone<br>- 4’,5,7-Trimethoxyflavone<br>- 5-Hydroxy-3,7,3’,4’-tetramethoxyflavone<br>- 8-Prenylnaringenin<br>- Acacetin<br>- Acacetin 3,6-di-C-glucoside<br>- Acacetin 8-C-neohesperidoside<br>- Acacetin 6-C-neohesperidoside<br>- Anthocyanidin<br>- Anthocyanin<br>- Apigenin<br>- Apigetrin<br>- Apigenin 8-C-neohesperidoside<br>- Catechin<br>- Cosmosiin<br>- Cosmosiine<br>- Cosmosioside<br>- Cosmetin<br>- Cosmosin<br>- Chalcone<br>- Chrysoeriol<br>- Chrysoeriol 7-O-neohesperidoside-4’-glucoside<br>- Chrysoeriol 7-O-neohesperidoside<br>- Didymin<br>- Diosmetin<br>- Diosmin<br>- Dihydrokaempferol<br>- Dihydroquercetin<br>- Epicatechin<br>- Eriocitrin<br>- Eriodictyol<br>- Fortunellin<br>- Hesperetin<br>- Hesperidin<br>- Heridictyol<br>- Heptamethoxy-flavone<br>- Isoscutellarein<br>- Isosakuranetin<br>- Isosinensetin<br>- Isovitexin<br>- Isorhoifolin<br>- Isosakuranetin 7-O-rutinoside<br>- Isosakuranetin 7-O-neohesperidoside<br>- Kämpferol<br>- Leucoanthocyanidin<br>- Luteolin<br>- Luteolin 7-O-rutinoside |

| | | |
|---|---|---|
| | | - Lucenin-2<br>- Lucenin-2 4’-methyl ether<br>- Natsudaidain<br>- Narirutin<br>- Naringenin<br>- Naringin<br>- Neodiosmin<br>- Neoeriocitrin<br>- Neohesperidin<br>- Nohesperidose<br>- Neoponcirin<br>- Nobiletin<br>- Orientin 4’-methyl ether<br>- Poncirin<br>- Pentamethylquercetin<br>- Quercetin<br>- Quercetogetin<br>- Rhioflin<br>- Rhoifolin 4’-glucoside<br>- Rutin<br>- Rutinose<br>- Scoparin<br>- Sinensetin<br>- Stellarin-2<br>- Sudachitin<br>- Tangeretin<br>- Thalictiin<br>- Taxifolin<br>- Tetramethylscutellarein<br>- Vicenin-2<br>- Xanthohumol<br><br>Exclusion criteria: Interventions that are not citrus flavonoids from the list above, or administered before model induction. |
| 26. | Outcome measures | Inclusion criteria:<br>Thermal behavioural responses:-<br>Evoked Limb Withdrawal (ELW) Heat –<br>Radiant heat (Hargreaves’)<br>Hot plate<br>Paw immersion<br>Evoked Limb Withdrawal (ELW) Cold –<br>Acetone<br>Cold probe<br>Cold plate<br>Cold tolerance<br>Limb immersion<br>Evoked Tail Withdrawal (ETW) Heat –<br>Tail flick<br>Tail immersion<br>Evoked Tail Withdrawal (ETW) Cold –<br>Tail immersion |

| | | |
|---|---|---|
| | | Exclusion criteria: Any outcomes that are not thermal-related pain behavioural outcomes |
| 27. | Language restrictions | Inclusion criteria: All languages<br>Exclusion criteria: None |
| 28. | Publication date restrictions | Inclusion criteria: All publication dates<br>Exclusion criteria: None |
| 29. | Other | Inclusion criteria:<br>Exclusion criteria: |
| 30. | Sort and prioritize your exclusion criteria per selection phase | Selection phase: *full text screening<br>1. All previously stated inclusion/exclusion criteria will be applied in this phase.<br><br>*most studies do not report exact outcomes in title/abstract so full text screening and data extraction are being carried out in one phase. |
| | Study characteristics to be extracted (for assessment of external validity, reporting quality) | |
| 31. | Study ID (*e.g.* authors, year) | - Title<br>- Primary Author<br>- Corresponding authors<br>- DOI<br>- Year<br>- Journal name<br>- Country of origin of corresponding author |
| 32. | Study design characteristics (*e.g.* experimental groups, number of animals) | Animal husbandry<br>- Number of animals per cage<br>- Light cycle of animal facilities<br>- Environmental enrichment reported<br>- Habituation time (days) |
| 33. | Animal model characteristics (*e.g.* species, gender, disease induction) | - Animal species<br>- Animal background strain<br>- Sex of animals<br>- Age of animals<br>- STZ dose for induction<br>- STZ route of administration |
| 34. | Intervention characteristics (*e.g.* intervention, timing, duration) | - Dose<br>- Timing<br>- Duration<br>- Route of administration |
| 35. | Outcome measures | - Outcome measure method<br>- Site of outcome measurement<br>- Number of animals per group<br>- Temperature (for thermal outcomes)<br>- Time (e.g. for paw withdrawal)<br>- Outcome measure units (e.g. sec for time)<br>- Outcome measure average type<br>- Outcome measure error type (SD/SEM)<br>- Point of data extraction (c) |
| 36. | Other (*e.g.* drop-outs) | |
| | Assessment risk of bias (internal validity) or study quality | |

| | | |
|---|---|---|
| 37. | Specify (a) the number of reviewers assessing the risk of bias/study quality in each study and (b) how discrepancies will be resolved | 2 reviewers will assess the risk of bias/study quality in each study.<br>Discrepancies resolved with a third expert party (Dr. Gillian Currie) |
| 38. | Define criteria to assess (a) the internal validity of included studies (*e.g.* selection, performance, detection and attrition bias) and/or (b) other study quality measures (*e.g.* reporting quality, power) | ☐By use of SYRCLE's Risk of Bias tool[4]<br>☐By use of SYRCLE's Risk of Bias tool, adapted as follows:<br>☐By use of CAMARADES' study quality checklist, e.g [22]<br>☒By use of CAMARADES' study quality checklist, adapted as follows:<br>- Reporting of random allocation of animals to experimental groups<br>- Reporting of blinding during outcome assessment<br>- Reporting of a sample size calculation<br>- Reporting of a st udy protocol dated before the study before the study began<br>- Reporting of rationale for data point inclusion/exclusion<br>- Reporting of potential conflicts of interest statement<br>- Reporting whether experiments were approved by an animal welfare committee<br>☐Other criteria, namely: |
| | Collection of outcome data | |
| 39. | For each outcome measure, define the type of data to be extracted (*e.g.* continuous/dichotomous, unit of measurement) | Continuous for all numerical variables<br>Dichotomous for categorical variables |
| 40. | Methods for data extraction/retrieval (*e.g.* first extraction from graphs using a digital screen ruler, then contacting authors) | Extraction of numerical data (mean, SD, or SEM) from the full texts of included papers using WebPlotDigitizer |
| 41. | Specify (a) the number of reviewers extracting data and (b) how discrepancies will be resolved | Two reviewers will extract data from each included publication and resolve discrepancies by discussion/involvement of third expert party (Dr. Gillian Currie) |
| | Data analysis/synthesis | |
| 42. | Specify (per outcome measure) how you are planning to combine/compare the data (*e.g.* descriptive summary, meta-analysis) | Findings from SR will be collated in a descriptive summary, paired with a meta-analysis of numerical data, if sufficient number of comparisons ($n \geq 10$) included. |
| 43. | Specify (per outcome measure) how it will be decided whether a meta-analysis will be performed | All numerical data will be included in a meta-analysis, with more weight given to studies with higher precision |
| | *If a meta-analysis seems feasible/sensible, specify (for each outcome measure):* | |
| 44. | The effect measure to be used (*e.g.* mean difference, standardized mean difference, risk ratio, odds ratio) | Standardised mean difference |

| | | |
|---|---|---|
| 45. | The statistical model of analysis (*e.g.* random or fixed effects model) | Multivariable meta-regression, if sufficient number of comparisons (n≥10) included. Alternatively, each characteristic will be investigated using a univariate approach. |
| 46. | The statistical methods to assess heterogeneity (*e.g.* $I^2$, Q) | Cochran's, $Tau^2$, residual $I^2$ |
| 47. | Which study characteristics will be examined as potential source of heterogeneity (subgroup analysis) | Study quality measures<br>- Reporting of random allocation of animals to experimental groups<br>- Reporting of blinding during outcome assessment<br>- Reporting of a sample size calculation<br>- Reporting of a study protocol dated before the study before the study began<br>- Reporting of rationale for data point inclusion/exclusion<br>Type of citrus flavonoids<br>Sex of animals |
| 48. | Any sensitivity analyses you propose to perform | Sensitivity analysis in R |
| 49. | Other details meta-analysis (*e.g.* correction for multiple testing, correction for multiple use of control group) | Where an intervention control serves more than one intervention, the number of animals in the control group would be corrected (Vesterinen et al, 2016).<br><br>Where several measurements of the same underlying biological outcome are reported from the same cohort of animals at the same time point, we will nest these comparisons into a single fixed effect (Vesterinen et al, 2016). |
| 50. | The method for assessment of publication bias | Funnel plot, egger's and trim and fill will be used to assess publication bias for each outcome (requires at least 25 comparisons) |
| Final approval by (names, affiliations): | | Date: |

**References**

1. Li, C., Bunner, A.E. and Pippin, J.J., 2013. From animal models to clinical practicality: lessons learned from current translational progress of diabetic peripheral neuropathy. Peripheral Neuropathy—A New Insight into the Mechanism, Evaluation and Management of a Complex Disorder. Rijeka, Hungary: InTech, pp.29-76.
2. IndINeP Study Group, 2008. Burden of neuropathic pain in Indian patients attending urban, specialty clinics: results from a cross sectional study. Pain Practice, 8(5), pp.362-378.
3. Van Acker, K., Bouhassira, D., De Bacquer, D., Weiss, S., Matthys, K., Raemen, H., Mathieu, C. and Colin, I.M., 2009. Prevalence and impact on quality of life of peripheral neuropathy with or without neuropathic pain in type 1 and type 2 diabetic patients attending hospital outpatients clinics. Diabetes & metabolism, 35(3), pp.206-213.

4. Feldman, E.L., Callaghan, B.C., Pop-Busui, R., Zochodne, D.W., Wright, D.E., Bennett, D.L., Bril, V., Russell, J.W. and Viswanathan, V., 2019. Diabetic neuropathy. Nature reviews Disease primers, 5(1), p.41.
5. Courteix, C., Eschalier, A. and Lavarenne, J., 1993. Streptozocin-induced diabetic rats: behavioural evidence for a model of chronic pain. Pain, 53(1), pp.81-88.
6. Junod, A.L.A.E.O.L.P.R.C.A.E.R.A.E., Lambert, A.E., Orci, L., Pictet, R., Gonet, A.E. and Renold, A.E., 1967. Studies of the diabetogenic action of streptozotocin. Proceedings of the Society for Experimental Biology and Medicine, 126(1), pp.201-205.
7. Ghasemi, A. and Jeddi, S., 2023. Streptozotocin as a tool for induction of rat models of diabetes: A practical guide. EXCLI journal, 22, p.274.
8. Mahmoud, A.M., Hernandez Bautista, R.J., Sandhu, M.A. and Hussein, O.E., 2019. Beneficial effects of citrus flavonoids on cardiovascular and metabolic health. Oxidative medicine and cellular longevity, 2019.
9. Li, C. and Schluesener, H., 2017. Health-promoting effects of the citrus flavanone hesperidin. Critical reviews in food science and nutrition, 57(3), pp.613-631.
10. Millar, C.L., Duclos, Q. and Blesso, C.N., 2017. Effects of dietary flavonoids on reverse cholesterol transport, HDL metabolism, and HDL function. Advances in Nutrition, 8(2), pp.226-239.
11. Rees, A., Dodd, G.F. and Spencer, J.P., 2018. The effects of flavonoids on cardiovascular health: A review of human intervention trials and implications for cerebrovascular function. Nutrients, 10(12), p.1852.
12. Zaidun, N.H., Thent, Z.C. and Abd Latiff, A., 2018. Combating oxidative stress disorders with citrus flavonoid: Naringenin. Life sciences, 208, pp.111-122.
13. Zhang, X., Li, X., Fang, H., Guo, F., Li, F., Chen, A. and Huang, S., 2019. Flavonoids as inducers of white adipose tissue browning and thermogenesis: signalling pathways and molecular triggers. Nutrition & metabolism, 16, pp.1-15.
14. Kopustinskiene, D.M., Jakstas, V., Savickas, A. and Bernatoniene, J., 2020. Flavonoids as anticancer agents. Nutrients, 12(2), p.457.
15. Vesterinen, H.M., Sena, E.S., Egan, K.J., Hirst, T.C., Churolov, L., Currie, G.L., Antonic, A., Howells, D.W. and Macleod, M.R., 2014. Meta-analysis of data from animal studies: a practical guide. Journal of neuroscience methods, 221, pp.92-102.
