## Supplementary material for "Machine-assisted systematic review and meta-analysis of citrus flavonoids in streptozotocin-induced painful diabetic neuropathy": Mechanical outcomes summary

### Mechanical Outcome Findings

| Paper | Citrus Flavonoid Tested | Dose (mg/kg) | Behavioural Test | Behavioural Test Measurement | Behavioural Test Measurement Result | Time of Outcome Measurement (weeks post citrus flavonoid administration) | Reduction of Pain-Related Behaviours |
|---|---|---|---|---|---|---|---|
| [1] | Naringin | 10 | Plantar Test | Paw Withdrawal Latency | 4.83 seconds +/- 0.75 | 4 | Yes |
| [1] | Naringin | 20 | Plantar Test | Paw Withdrawal Latency | 4.33 seconds +/- 0.816 | 4 | Yes |
| [1] | Naringin | 40 | Plantar Test | Paw Withdrawal Latency | 2.66 seconds +/- 1.21 | 4 | Yes |
| [1] | Naringin | Diabetic Control | Plantar Test | Paw Withdrawal Latency | 5.16 seconds +/- 0.75 | 4 | [redacted] |
| [1] | Naringin | Naïve Control | Plantar Test | Paw Withdrawal Latency | 2.33 seconds +/- 1.36 | 4 | [redacted] |
| [2] | Naringin | 20 | Von Frey | Paw Withdrawal Threshold | 29.934g +/- 1.59 | 4 | Yes |
| [2] | Naringin | 40 | Von Frey | Paw Withdrawal Threshold | 47.152g +/- 2.252 | 4 | Yes |
| Y | Naringin | 80 | Von Frey | Paw Withdrawal Threshold | 53.775g +/- 1.722 | 4 | Yes |
| [2] | Naringin | Diabetic Control | Von Frey | Paw Withdrawal Threshold | 13.775g +/- 2.517 | 4 | [redacted] |
| [2] | Naringin | Naïve Control | Von Frey | Paw Withdrawal Threshold | 73.642g +/- 2.384 | 4 | [redacted] |
| [2] | Naringin | 20 | Randall-Selitto | Paw Withdrawal Threshold | 87.5g +/- 10.577 | 4 | No |
| [2] | Naringin | 40 | Randall-Selitto | Paw Withdrawal Threshold | 131.731g +/- 12.5 | 4 | Yes |
| [2] | Naringin | 80 | Randall-Selitto | Paw Withdrawal Threshold | 184.615g +/- 6.731 | 4 | Yes |
| [2] | Naringin | Diabetic Control | Randall-Selitto | Paw Withdrawal Threshold | 50g +/- 9.615 | 4 | [redacted] |
| [2] | Naringin | Naïve Control | Randall-Selitto | Paw Withdrawal Threshold | 267.308g +/- 9.616 | 4 | [redacted] |
| [3] | Naringenin | 10 | Von Frey | Paw Withdrawal Threshold | Not recorded | Not Recorded | [redacted] |
| [3] | Naringenin | 20 | Von Frey | Paw Withdrawal Threshold | 6.007g +/- 0.878 | 8 | No |
| [3] | Naringenin | 50 | Von Frey | Paw Withdrawal | 12.580g +/- 0.476 | 8 | Yes |

| | | | | | | | |
|---|---|---|---|---|---|---|---|
| | | | | Threshold | | | |
| [3] | Naringenin | 100 | Von Frey | Paw Withdrawal Threshold | 12.580g +/- 0.892 | 8 | Yes |
| [3] | Naringenin | Diabetes Control | Von Frey | Paw Withdrawal Threshold | 3.896g +/- 0.684 | 8 | |
| [3] | Naringenin | Naïve Control | Von Frey | Paw Withdrawal Threshold | 14.840g +/- 0.179 | 8 | |
| [3] | Naringenin | Naïve 10mg/kg | Von Frey | Paw Withdrawal Threshold | Not recorded | Not Recorded | |
| [3] | Naringenin | Naïve 20mg/kg | Von Frey | Paw Withdrawal Threshold | 12.729g +/- 0.313 | 8 | |
| [3] | Naringenin | Naïve 50mg/kg | Von Frey | Paw Withdrawal Threshold | 13.680g +/- 0.491 | 8 | |
| [3] | Naringenin | Naïve 100mg/kg | Von Frey | Paw Withdrawal Threshold | 12.996g +/- 0.536 | 8 | |
| [4] | Naringenin | 25 | Randall-Selitto | Paw-Withdrawal Pressure | 295.597mmHg +/- 21.227 | 5 | No |
| [4] | Naringenin | 50 | Randall-Selitto | Paw-Withdrawal Pressure | 339.623mmHg +/- 18.082 | 5 | Yes |
| [4] | Naringenin | Diabetic Control | Randall-Selitto | Paw-Withdrawal Pressure | 257.862mmHg +/- 17.296 | 5 | |
| [4] | Naringenin | Naïve Control | Randall-Selitto | Paw-Withdrawal Pressure | 367.925mmHg +/- 22.013 | 5 | |
| [5] | Naringenin | 25 | Randall-Selitto - left paw | Pain Threshold | 97.17g +/- 1.887 | 4 | Yes |
| [5] | Naringenin | 50 | Randall-Selitto - left paw | Pain Threshold | 97.17g +/- 4.717 | 4 | Yes |
| [5] | Naringenin | 100 | Randall-Selitto - left paw | Pain Threshold | 85.849g +/- 1.887 | 4 | Yes |
| [5] | Naringenin | Diabetic Control | Randall-Selitto - left paw | Pain Threshold | 98.113g +/- 3.774 | 4 | |
| [5] | Naringenin | Naïve Control | Randall-Selitto - left paw | Pain Threshold | 171.698g +/- 9.434 | 4 | |
| [5] | Naringenin | Naïve + 50mg/kg | Randall-Selitto - left paw | Pain Threshold | 161.321g +/- 2.83 | 4 | |
| [5] | Naringenin | 25 | Randall-Selitto - right paw | Pain Threshold | 95.874g +/- 3.641 | 4 | Yes |
| [5] | Naringenin | 50 | Randall-Selitto - right paw | Pain Threshold | 86.165g +/- 6.068 | 4 | Yes |
| [5] | Naringenin | 100 | Randall-Selitto - right paw | Pain Threshold | 98.301g +/- 4.854 | 4 | Yes |
| [5] | Naringenin | Diabetic Control | Randall-Selitto - right | Pain Threshold | 91.019g +/- 6.068 | 4 | |

| | | | | | | | |
|---|---|---|---|---|---|---|---|
| | | | paw | | | | |
| [5] | Naringenin | Naïve Control | Randall-Selitto - right paw | Pain Threshold | 192.961g +/- 4.855 | 4 | |
| [5] | Naringenin | Naïve + 50mg/kg | Randall-Selitto - right paw | Pain Threshold | 169.903g +/- 1.820 | 4 | |
| [5] | Naringenin | 25 | Von Frey - left paw | Pain Threshold | 34.421g +/- 1.891 | 4 | No |
| [5] | Naringenin | 50 | Von Frey - left paw | Pain Threshold | 39.338g +/- 4.728 | 4 | Yes |
| [5] | Naringenin | 100 | Von Frey - left paw | Pain Threshold | 31.206 +/- 0.945 | 4 | Yes |
| [5] | Naringenin | Diabetic Control | Von Frey - left paw | Pain Threshold | 34.61g +/- 1.513 | 4 | |
| [5] | Naringenin | Naïve Control | Von Frey - left paw | Pain Threshold | 67.707g +/- 1.324 | 4 | |
| [5] | Naringenin | Naïve + 50 | Von Frey - left paw | Pain Threshold | 71.3g +/- 0.757 | 4 | |
| [5] | Naringenin | 25 | Von Frey - right paw | Pain Threshold | 31.538g +/- 1.347 | 4 | No |
| [5] | Naringenin | 50 | Von Frey - right paw | Pain Threshold | 39.038g +/- 1.731 | 4 | Yes |
| [5] | Naringenin | 100 | Von Frey - right paw | Pain Threshold | 35.769 +/- 2.308 | 4 | Yes |
| [5] | Naringenin | Diabetic Control | Von Frey - right paw | Pain Threshold | 38.077g +/- 1.538 | 4 | |
| [5] | Naringenin | Naïve Control | Von Frey - right paw | Pain Threshold | 65.385g +/- 1.538 | 4 | |
| [5] | Naringenin | Naïve + 50mg/kg | Von Frey - right paw | Pain Threshold | 66.346g +/- 1.154 | 4 | |
| [6] | Rutin | 100 | Von Frey | Paw Withdrawal Threshold | 60.526g +/- 0.878 | 8 | Yes |
| [6] | Rutin | 200 | Von Frey | Paw Withdrawal Threshold | 65.497g +/- 1.462 | 8 | Yes |
| [6] | Rutin | Diabetic Control | Von Frey | Paw Withdrawal Threshold | 33.431g +/- 0.880 | 8 | |
| [6] | Rutin | Naive Control | Von Frey | Paw Withdrawal Threshold | 73.9g +/- 1.467 | 8 | |
| [6] | Rutin | 100 | Randall-Selitto | Paw Withdrawal Threshold | 133.632g +/- 1.918 | 8 | Yes |
| [6] | Rutin | 200 | Randall-Selitto | Paw Withdrawal Threshold | 138.107g +/- 1.919 | 8 | Yes |
| [6] | Rutin | Diabetic Control | Randall-Selitto | Paw Withdrawal Threshold | 67.136g +/- 2.557 | 8 | |
| [6] | Rutin | Naive Control | Randall-Selitto | Paw Withdrawal Threshold | 187.98g +/- 0.819 | 8 | |
| [7] | Quercetin | 10 | Randall-Selitto | Paw Withdrawal Threshold | 118.1g +/- 12.884 | 8 | No |
| [7] | Quercetin | 20 | Randall-Selitto | Paw Withdrawal Threshold | 157.822g +/- 11.274 | 8 | Yes |
| [7] | Quercetin | 40 | Randall-Selitto | Paw Withdrawal | 199.693g +/- 9.126 | 8 | Yes |

| | | | | | | | |
|---|---|---|---|---|---|---|---|
| | | | | Threshold | | | |
| [7] | Quercetin | Diabetic Control | Randall-Selitto | Paw Withdrawal Threshold | 83.742g +/- 11.81 | 8 | |
| [7] | Quercetin | Naive Control | Randall-Selitto | Paw Withdrawal Threshold | 280.217g +/- 11.81 | 8 | |
| [7] | Quercetin | Naive + 40mg/kg | Randall-Selitto | Paw Withdrawal Threshold | 283.44g +/- 11.809 | 8 | |
| [7] | Quercetin | 10 | Von Frey | Paw Withdrawal Threshold | 37.374g +/- 3.031 | 8 | No |
| [7] | Quercetin | 20 | Von Frey | Paw Withdrawal Threshold | 55.892g +/- 3.872 | 8 | Yes |
| [7] | Quercetin | 40 | Von Frey | Paw Withdrawal Threshold | 69.024g +/- 4.209 | 8 | Yes |
| [7] | Quercetin | Diabetic Control | Von Frey | Paw Withdrawal Threshold | 23.569g +/- 2.189 | 8 | |
| [7] | Quercetin | Naive Control | Von Frey | Paw Withdrawal Threshold | 95.286g +/- 4.377 | 8 | |
| [7] | Quercetin | Naive + 40mg/kg | Von Frey | Paw Withdrawal Threshold | 90.236g +/- 5.051 | 8 | |
| [8] | Hesperidin | 25 | Randall-Selitto | Paw Withdrawal Threshold | 101.559g +/- 8.686 | 4 | Yes |
| [8] | Hesperidin | 50 | Randall-Selitto | Paw Withdrawal Threshold | 131.626g +/- 12.027 | 4 | Yes |
| [8] | Hesperidin | 100 | Randall-Selitto | Paw Withdrawal Threshold | 165.033g +/- 10.691 | 4 | Yes |
| [8] | Hesperidin | Diabetic Control | Randall-Selitto | Paw Withdrawal Threshold | 50.78g +/- 8.018 | 4 | |
| [8] | Hesperidin | Naive Control | Randall-Selitto | Paw Withdrawal Threshold | 281.96g +/- 12.695 | 4 | |
| [8] | Hesperidin | 25 | Von Frey | Paw Withdrawal Threshold | 30.145g +/- 2.705 | 4 | Yes |
| [8] | Hesperidin | 50 | Von Frey | Paw Withdrawal Threshold | 40.580g +/- 2.513 | 4 | Yes |
| [8] | Hesperidin | 100 | Von Frey | Paw Withdrawal Threshold | 51.401g +/- 3.092 | 4 | Yes |
| [8] | Hesperidin | Diabetic Control | Von Frey | Paw Withdrawal Threshold | 17.391g +/- 3.285 | 4 | |
| [8] | Hesperidin | Naive Control | Von Frey | Paw Withdrawal Threshold | 80.773g +/- 5.024 | 4 | |
| [9] | Luteolin | 50 | Von frey | Mechanical Withdrawal Threshold | 7.404g +/- 0.575 | 3 | Yes |
| [9] | Luteolin | 100 | Von frey | Mechanical Withdrawal | 9.511g +/- 0.893 | 3 | Yes |

| | | | | | | | |
|---|---|---|---|---|---|---|---|
| | | | | Threshold | | | |
| [9] | Luteolin | 200 | Von frey | Mechanical Withdrawal Threshold | 9.128g +/- 0.830 | 3 | Yes |
| [9] | Luteolin | Diabetic Control | Von frey | Mechanical Withdrawal Threshold | 4.277g +/- 0.511 | 3 | |
| [9] | Luteolin | Naive Control | Von frey | Mechanical Withdrawal Threshold | 11.298g +/- 1.214 | 3 | |
