## Supplementary material for "Machine-assisted systematic review and meta-analysis of citrus flavonoids in streptozotocin-induced painful diabetic neuropathy": R script for Meta Analysis

### R code for meta-analysis

```
library(tidyverse)
library(rmarkdown)
library(meta)
library(metafor)
library(readxl)
library(dplyr)
library(stringr)


# read in excel file (SyRF output or modified SyRF output)
thermal_outcomes <- read_excel("Downloads/Quantitative Syrf Data Shreya.xlsx")

modelling_data_interventions <- thermal_outcomes %>%
  group_by(StudyId, ExperimentId) %>%
  filter(any(IsTreatmentControl == TRUE) &
         any(IsTreatmentControl == FALSE)) %>%
  filter(!(DiseaseModelLabel == "Naive")) %>%
  filter(!is.na(IsTreatmentControl))

#DATA CLEANING

# Condense the three strain columns into one, replacing "Other" with specific strains
modelling_data_interventions <- modelling_data_interventions %>%
  mutate(ConsolidatedStrain = case_when(
    Strain != "Other" ~ Strain,
    Strain == "Other" & !is.na(`State other strain`) ~ `State other strain`,
    Strain == "Other" & is.na(`State other strain`) ~ `Strain of mouse`,
    TRUE ~ `Strain of mouse`
  ))

# Condense the "Route of Administration" and "If other, state route" columns into one
modelling_data_interventions <- modelling_data_interventions %>%
  mutate(ConsolidatedRouteofIntervention = case_when(
    `Route of Administration` != "Other" ~ `Route of Administration`,
    `Route of Administration` == "Other" & !is.na(`If other, state route...76`) ~ `If other, state
route...76`,
    TRUE ~ as.character(`Route of Administration`)
  ))


# Standardizing "Tail-flick" and "Tail flick" to a single format
modelling_data_interventions <- modelling_data_interventions %>%
```

```
  mutate(OutcomeLabel = str_replace_all(OutcomeLabel, fixed("Tail flick"), "Tail-flick"))

# Standardizing "Wistar-Kyoto" and "Wistar Albino" to a single format
modelling_data_interventions <- modelling_data_interventions %>%
  mutate(ConsolidatedStrain = str_replace(ConsolidatedStrain, "Wistar Albino",
"Wistar")) %>%
  mutate(ConsolidatedStrain = str_replace(ConsolidatedStrain, "Wistar-Kyoto", "Wistar"))



# Select relevant study characteristics and characteristics needed for meta-analysis
(columns in SyRF)
modelling_data_interventions <- modelling_data_interventions %>%
  select(StudyId, ExperimentId, CohortId, DiseaseModelLabel, DiseaseModelId,
NumberOfAnimals, OutcomeId, GreaterIsWorse, OutcomeAverage, OutcomeError,
OutcomeAverageType, ErrorType, `Sex of animals`, `Weight of animals`, `Was Random
Allocation To Group Reported`, Species, `Was Allocation Concealment Reported?`, `Was
Blinding Used During Outcome Assessment?`, `Did the authors report using a sample size
calculation?`,
        `Were Any Experimental Animals Excluded From Analysis?`, `Do the authors report
that the study was pre-registered?`, `Is Compliance with Animal Welfare Regulations
Reported?`, `Do the authors report a potential conflict of interest statement?`, TimeInMinute,
        OutcomeLabel, TreatmentId, TreatmentLabel, `Timing of Administration of Intervention
post STZ induction (in days)`, IsTreatmentControl, ConsolidatedRouteofIntervention,
ConsolidatedStrain, `Dose (mg/kg)`, `Number of Doses`, `Duration of Treatment (in weeks)`,
`Citrus Flavonoid Used`, `Was baseline pain measured pre-intervention admin?`, Authors,
Year, `STZ dose for diabetes induction`)

more_than_2_comparisons_interventions <- modelling_data_interventions %>%
  select(StudyId, ExperimentId, CohortId, IsTreatmentControl, OutcomeId) %>%
  group_by(StudyId, ExperimentId, OutcomeId) %>%
  dplyr::filter(IsTreatmentControl == TRUE) %>%
  count() %>%
  dplyr::filter(n>1) %>%
  unique()

# Make sure SD and SEM both exist/are calculated
modelling_data_final_interventions <- modelling_data_interventions %>%
  mutate(SEM = ifelse(ErrorType == "SEM", OutcomeError, NA)) %>%
  mutate(SD = ifelse(ErrorType == "SD", OutcomeError,
                  try(SEM * sqrt(NumberOfAnimals), silent=T))) %>%
  select(-OutcomeError) %>% # to be replaced with either SD or SEM
  mutate(Direction = ifelse(GreaterIsWorse ==TRUE, 1, -1)) #if greater is worse, make
direction of effect 1, else make it -1

# Example test names for classification - replace these with actual test names from your
dataset
```

```
heat_related_tests <- c("Hot plate", "Hargreaves"", "Tail-flick", "Tail immersion (warm water)")
# Example heat-related tests
cold_related_tests <- c("Cold plate", "Acetone drop", "Tail immersion (cold water)") #
Example cold-related tests

# Add a new column for outcome type classification based on the test name
modelling_data_final_interventions <- modelling_data_final_interventions %>%
  mutate(OutcomeType = case_when(
    OutcomeLabel %in% heat_related_tests ~ "Heat-related",
    OutcomeLabel %in% cold_related_tests ~ "Cold-related",
    TRUE ~ "Other"  # Use "Other" or NA for tests that don't fit either category
  ))

# Proceed with analysis using the classified data

# check numerical data is NUMERIC type (not character)
modelling_data_final_interventions <- modelling_data_final_interventions %>%
  mutate(NumberOfAnimals = as.numeric(NumberOfAnimals)) %>%
  mutate(OutcomeAverage = as.numeric(OutcomeAverage)) %>%
  mutate(SEM = as.numeric(SEM)) %>%
  mutate(`Timing of Administration of Intervention post STZ induction (in days)` =
as.numeric(`Timing of Administration of Intervention post STZ induction (in days)`)) %>%
  mutate(TimeInMinute = as.numeric(TimeInMinute)) %>%
  mutate(`Dose (mg/kg)` = as.numeric(`Dose (mg/kg)`)) %>%
  mutate(`STZ dose for diabetes induction` = as.numeric(`STZ dose for diabetes
induction`)) %>%
  mutate(`Number of Doses` = as.numeric(`Number of Doses`)) %>%
  mutate(`Duration of Treatment (in weeks)` = as.numeric(`Duration of Treatment (in
weeks)`)) %>%
  mutate(SD = as.numeric(SD))

# pivot to make data wide format - with experimental groups on same row
# Select values from variables where value could differ for intervention and control
# each line should then have a value for intervention and for control for each selected
variable

modelling_data_final_interventions <- modelling_data_final_interventions %>%
  pivot_wider(names_from = IsTreatmentControl, values_from=c(NumberOfAnimals, SD,
SEM, OutcomeAverage, DiseaseModelId, DiseaseModelLabel, CohortId, TreatmentId,
TreatmentLabel, ConsolidatedRouteofIntervention, `Dose (mg/kg)`, `Duration of Treatment
(in weeks)`, `Number of Doses`, `Citrus Flavonoid Used`, TimeInMinute, `Timing of
Administration of Intervention post STZ induction (in days)`))

# unnest data
tnc <- nrow(modelling_data_final_interventions)
for(i in 1:tnc){
```

```
  lsd <- modelling_data_final_interventions[i,27] %>% unnest(cols = everything() ) # choose
column number of animals in intervention group
  modelling_data_final_interventions[i,58] <- nrow(lsd) #first empty column <- choose column
count of number of model groups earlier combined into one
}
modelling_data_final_interventions <- modelling_data_final_interventions %>%
  unnest(cols = everything() )

tnd <- nrow(modelling_data_final_interventions)

# Calculate true number of control animals, i.e. N/number of comparisons served by control
group
trueC <- data.frame()
for(i in 1:tnd){
  trueC[i,1] <-
(modelling_data_final_interventions[i,26]/modelling_data_final_interventions[i,58]) #number
of animals in control group divided by number of model groups it serves
}
modelling_data_final_interventions <- cbind(modelling_data_final_interventions, trueC)

# calculate SMD effect sizes, standard errors of SMD, weights, and pooled SD (from
Vesterinen et al., 2014)
modelling_data_final_interventions <- modelling_data_final_interventions %>%
  mutate(combined_n = `NumberOfAnimals_TRUE...26` +
`NumberOfAnimals_FALSE`) %>% # N control + N disease model
  mutate(SPooled = sqrt(((`NumberOfAnimals_TRUE...26` - 1) * `SD_TRUE`^2 +
                         (NumberOfAnimals_FALSE - 1) * SD_FALSE^2) / (combined_n-2))) %>%
#equation 6 from paper
  mutate(SMD_ES = ((abs(`OutcomeAverage_TRUE`)-
abs(`OutcomeAverage_FALSE`))/SPooled) * # standardised mean difference
           (1 - 3/(4*combined_n-9)) * Direction) %>% #equation 15
  mutate(SMD_ES_SE  = sqrt((combined_n/(NumberOfAnimals_FALSE *
`NumberOfAnimals_TRUE...26`)) + # standard error
                           SMD_ES^2/(2*(combined_n - 3.94)))) %>% #equation 16
  mutate(weight = 1/(SMD_ES_SE^2)) %>% # weight studies by inverse error squared
  mutate(weighted_ES = SMD_ES * weight) %>%
  filter(!is.na(weighted_ES)) # remove the few rows where effect size calculation wasn't
possible (where there wasn't a model and control group)



# if you want to calculate a nested effect size and error (where several effect sizes are
related they can be combined into a single estimate)
# MA Code nesting - one effect size for each cohort

Nested_all_1 <- modelling_data_final_interventions %>%
```

```
  group_by(StudyId, ExperimentId, TreatmentId_FALSE) %>% # group by study Id and by
experiment ID (if one control by experiment)
  mutate(Nested_ES = sum(weighted_ES)/sum(weight)) %>% # calculated nested effect size
  mutate(NestN = length(OutcomeAverage_FALSE)) %>% # number of studies to nest
  mutate(Nested_ES_SE = ifelse(NestN>1, sqrt(NestN/sum(weight)),SMD_ES_SE)) %>% #
if more than one effect size to nest, calculate nested value, otherwise equal to SMD_SE
from earlier
  dplyr::select(StudyId, ExperimentId, OutcomeAverageType, ErrorType, `Sex of animals`,
`Weight of animals`, `Was Random Allocation To Group Reported`, Species, `Was Allocation
Concealment Reported?`, `Was Blinding Used During Outcome Assessment?`, `Did the
authors report using a sample size calculation?`,
              `Were Any Experimental Animals Excluded From Analysis?`, `Do the authors
report that the study was pre-registered?`, `Is Compliance with Animal Welfare Regulations
Reported?`, `Do the authors report a potential conflict of interest statement?`,
              ConsolidatedStrain, `Was baseline pain measured pre-intervention admin?`,
Authors, Year,
              Nested_ES, NestN, Nested_ES_SE, TreatmentId_FALSE,
TreatmentLabel_FALSE, `Citrus Flavonoid Used_FALSE`, `Timing of Administration of
Intervention post STZ induction (in days)_FALSE`, `STZ dose for diabetes induction`,
`Duration of Treatment (in weeks)_FALSE`) %>%
  unique() %>%
  filter(!is.na(Nested_ES_SE))

# Build a new random-effects model
Nested_all_1_MA <- metagen(
  `Nested_ES`,
  `Nested_ES_SE`,
  sm = "SMD",
  prediction = TRUE,
  data = Nested_all_1,
  studlab = StudyId,
  random = TRUE,
  fixed = FALSE,
  method.tau = "REML",
)

Nested_all_1_MA

# Split the Authors column to extract the first author
# Assume Authors column is a string with authors separated by commas
Nested_all_1$FirstAuthor <- sapply(strsplit(Nested_all_1$Authors, "[,;]"), `[`, 1)

# Combine first author and year into a new column
Nested_all_1$StudyLabel <- paste(Nested_all_1$FirstAuthor, Nested_all_1$Year,
Nested_all_1$TreatmentLabel_FALSE, sep=", ")
```

```
# Extract the necessary data for the plot
study_labels <- Nested_all_1$StudyLabel

Q_value <- Nested_all_1_MA$Q
p_value <- Nested_all_1_MA$'p-value'

# Set plot dimensions and font size
plot_width <- 900
plot_height <- 800
base_font_size <- 12

# Specify colors for different elements
col_square <- "#56B4E9" # Light blue color for study squares
col_diamond <- "#D55E00" # Orange color for summary diamond
col_predict <- "#009E73" # Teal color for prediction interval

# Open PNG device
png(filename="Nested outcomes per cohort GPT 2.png", width=plot_width,
height=plot_height)

# Create forest plot
forest_plot_nested_all <- forest(
  Nested_all_1_MA,
  sortvar = TE,
  comb.fixed = FALSE,
  comb.random = TRUE,
  text.random=TRUE,
  prediction = TRUE,
  xlim=c(-5,10),
  studlab = study_labels, # specify the study labels here
  rightcols = c("effect", "ci"), # Adjust as needed to display effect sizes and CIs
  leftcols = "studlab",  # Adjust to display the study labels on the left
  xlab = "Standardized Mean Difference (SMD)", # Add x-axis label
  smlab = "SMD",
  label.right = "Improved Pain",
  label.left = "Worsened Pain",
  fontsize = base_font_size,
  plotwidth = paste0(plot_width/100, "cm"), # Convert pixels to cm
  plotheight = paste0(plot_height/100, "cm"), # Convert pixels to cm
  digits = 4,
  digits.se = 4,
  col.study = col_square,
  col.diamond = col_diamond,
  col.predict = col_predict,
  col.inside = "black", # Color for lines representing CIs
  cex = 1.5, # Enlargement factor for all text elements
```

```
  cex.axis = 0.7, # Enlargement factor for axis annotations
  cex.lab = 0.7, # Enlargement factor for axis titles
  colgap.fac = 0.5, # Factor for the gap between columns
  boxsize = 0.5, # Size of the squares representing individual studies
  pch = 15, # Symbol used for the point estimates in the forest plot
  pch.random = 15 # Symbol used for the pooled estimate in random effects model
)

text(x, y, labels=paste("Q =", formatC(Q_value, format='f', digits=2),
                        "p =", formatC(p_value, format='f', digits=4)))


dev.off()




################## HETEROGENEITY ANALYSIS ##########################

# STUDY DESIGN

#Flavonoid

Flavonoid <- Nested_all_1 %>%
  mutate(`Citrus Flavonoid Used_FALSE` = as.factor(`Citrus Flavonoid Used_FALSE`)) #
make sure variable is a character or factor#replace missing



Flavonoid_MA <- metagen(
  `Nested_ES`,
  `Nested_ES_SE`,
  sm = "SMD",
  prediction = TRUE,
  data = Flavonoid,
  subgroup = `Citrus Flavonoid Used_FALSE`, # add your subgroup here
  #tau.common = FALSE,
  studlab = StudyId,
  random = TRUE,
  fixed = FALSE,
  method.tau = "REML",
  #hakn = TRUE
)

png(file="Forest plot by flavonoid 2.png", width = 750, height =480)
```

```
forest(Flavonoid_MA, # use the meta object with your subgroup specified
       sortvar = TE,
       print.tau2 = FALSE,
       study.results = FALSE,
       subgroup = TRUE,
       label.right= "IMPROVED PAIN",
       label.left = "WORSENED PAIN",
       xlim=c(-7,10),
       layout="JAMA"
)

dev.off()



#Sex

Sex1 <- Nested_all_1 %>%
  mutate(`Sex of animals` = ifelse(is.na(`Sex of animals`),'NR',`Sex of animals`)) %>%
  mutate(`Sex of animals` = as.factor(`Sex of animals`)) # make sure variable is a character
or factor#replace missing

Sex1_MA <- metagen(
  `Nested_ES`,
  `Nested_ES_SE`,
  sm = "SMD",
  prediction = TRUE,
  data = Sex1,
  subgroup = `Sex of animals`, # add your subgroup here
  #tau.common = FALSE,
  studlab = StudyId,
  random = TRUE,
  fixed = FALSE,
  method.tau = "REML",
  #hakn = TRUE
)

png(file="Forest plot by sex 2.png", width = 770, height =350)

forest(Sex1_MA, # use the meta object with your subgroup specified
       sortvar = TE,
       print.tau2 = FALSE,
       study.results = FALSE,
       subgroup = TRUE,
       label.right= "IMPROVED PAIN",
       label.left = "WORSENED PAIN",
       xlim=c(-7,10),
```

```
        layout="JAMA",
)

dev.off()


 #STUDY QUALITY


# model without subgroup analysis
Nested_all_1_MA <- metagen(
  `Nested_ES`,
  `Nested_ES_SE`,
  sm = "SMD",
  prediction = TRUE,
  data = Nested_all_1,
  #tau.common = FALSE,
  studlab = StudyId,
  random = TRUE,
  fixed = FALSE,
  method.tau = "REML",
  #hakn = TRUE
)

#randomisation
Nested_all_1_MA <- metagen(
  `Nested_ES`,
  `Nested_ES_SE`,
  sm = "SMD",
  prediction = TRUE,
  data = Nested_all_1,
  subgroup = `Was Random Allocation To Group Reported`,
  #tau.common = FALSE,
  studlab = StudyId,
  random = TRUE,
  fixed = FALSE,
  method.tau = "REML",
  #hakn = TRUE
)

png(file="Forest plot randomisation 2.png", width = 770, height =350)

forest(Nested_all_1_MA,
       sortvar = TE,
       print.tau2 = FALSE,
       study.results = FALSE,
```

```
       subgroup = TRUE,
       label.right= "IMPROVED PAIN",
       label.left = "WORSENED PAIN",
       xlim=c(-5,6),
       layout="JAMA"
)

dev.off()

#blinding
Nested_all_1_MA <- metagen(
  `Nested_ES`,
  `Nested_ES_SE`,
  sm = "SMD",
  prediction = TRUE,
  data = Nested_all_1,
  subgroup = `Was Blinding Used During Outcome Assessment?`,
  #tau.common = FALSE,
  studlab = StudyId,
  random = TRUE,
  fixed = FALSE,
  method.tau = "REML",
  #hakn = TRUE
)

# plot subforest plot
png(file="Forest plot blinding 2.png", width = 770, height =350)

forest(Nested_all_1_MA,
       sortvar = TE,
       print.tau2 = FALSE,
       study.results = FALSE,
       subgroup = TRUE,
       label.right= "IMPROVED PAIN",
       label.left = "WORSENED PAIN",
       xlim=c(-5,6),
       layout="JAMA"
)

dev.off()


# POST-HOC

#Strain
```

```
Strain1 <- Nested_all_1 %>%
  mutate(ConsolidatedStrain = as.factor(ConsolidatedStrain)) # make sure variable is a
character or factor#replace missing



Strain1_MA <- metagen(
  `Nested_ES`,
  `Nested_ES_SE`,
  sm = "SMD",
  prediction = TRUE,
  data = Strain1,
  subgroup = ConsolidatedStrain, # add your subgroup here
  #tau.common = FALSE,
  studlab = StudyId,
  random = TRUE,
  fixed = FALSE,
  method.tau = "REML",
  #hakn = TRUE
)

png(file="Forest plot by strain 2.png", width = 750, height =400)

forest(Strain1_MA, # use the meta object with your subgroup specified
       sortvar = TE,
       print.tau2 = FALSE,
       study.results = FALSE,
       subgroup = TRUE,
       label.right= "IMPROVED PAIN",
       label.left = "WORSENED PAIN",
       xlim=c(-6,8),
       layout="JAMA"
)

dev.off()



#Baseline Pain Measurements

BaselinePain <- Nested_all_1 %>%
  mutate(`Was baseline pain measured pre-intervention admin?`
 = as.factor(`Was baseline pain measured pre-intervention admin?`)) # make sure variable is
a character or factor#replace missing



BaselinePain_MA <- metagen(
  `Nested_ES`,
```

```
  `Nested_ES_SE`,
  sm = "SMD",
  prediction = TRUE,
  data = BaselinePain,
  subgroup = `Was baseline pain measured pre-intervention admin?`
, # add your subgroup here
  #tau.common = FALSE,
  studlab = StudyId,
  random = TRUE,
  fixed = FALSE,
  method.tau = "REML",
  #hakn = TRUE
)

png(file="Forest plot by baseline pain 2.png", width = 750, height =350)

forest(BaselinePain_MA, # use the meta object with your subgroup specified
       sortvar = TE,
       print.tau2 = FALSE,
       study.results = FALSE,
       subgroup = TRUE,
       label.right= "IMPROVED PAIN",
       label.left = "WORSENED PAIN",
       xlim=c(-6,8),
       layout="JAMA"
)

dev.off()

#Outcome type - NOT ON NESTED DATA

TypeofOutcome <- modelling_data_final_interventions %>%
  mutate(OutcomeType = as.factor(OutcomeType)) # make sure variable is a character or
factor#replace missing



TypeofOutcome_MA <- metagen(
  `weighted_ES`,
  `SMD_ES_SE`,
  sm = "SMD",
  prediction = TRUE,
  data = TypeofOutcome,
  subgroup = OutcomeType, # add your subgroup here
  #tau.common = FALSE,
  studlab = StudyId,
  random = TRUE,
```

```
  fixed = FALSE,
  method.tau = "REML",
  #hakn = TRUE
)

png(file="Forest plot by outcome type 2.png", width = 750, height =350)

forest(TypeofOutcome_MA, # use the meta object with your subgroup specified
       sortvar = TE,
       print.tau2 = FALSE,
       study.results = FALSE,
       subgroup = TRUE,
       label.right= "IMPROVED PAIN",
       label.left = "WORSENED PAIN",
       xlim=c(-6,8),
       layout="JAMA"
)

dev.off()

#Pain test - NOT ON NESTED DATA

PainTest <- modelling_data_final_interventions %>%
  mutate(OutcomeLabel = as.factor(OutcomeLabel)) # make sure variable is a character or
factor#replace missing



PainTest_MA <- metagen(
  `weighted_ES`,
  `SMD_ES_SE`,
  sm = "SMD",
  prediction = TRUE,
  data = PainTest,
  subgroup = OutcomeLabel, # add your subgroup here
  #tau.common = FALSE,
  studlab = StudyId,
  random = TRUE,
  fixed = FALSE,
  method.tau = "REML",
  #hakn = TRUE
)

png(file="Forest plot by pain test 2.png", width = 750, height =550)

forest(PainTest_MA, # use the meta object with your subgroup specified
       sortvar = TE,
```

```
      print.tau2 = FALSE,
      study.results = FALSE,
      subgroup = TRUE,
      label.right= "IMPROVED PAIN",
      label.left = "WORSENED PAIN",
      xlim=c(-6,8),
      layout="JAMA"
)

dev.off()



#META REGRESSION

# Timing of intervention post STZ
# Example of renaming for simplicity
names(Nested_all_1)[names(Nested_all_1) == "Timing of Administration of Intervention post
STZ induction (in days)_FALSE"] <- "TimePostSTZ"

Nested_all_1_MA <- metagen(
  `Nested_ES`,
  `Nested_ES_SE`,
  sm = "SMD",
  prediction = TRUE,
  data = Nested_all_1,
  studlab = StudyId,
  random = TRUE,
  fixed = FALSE,
  method.tau = "REML",
)

Nested_all_1_MA

# Then, your formula would simply be
TimePostSTZ_metaregression <- metareg(Nested_all_1_MA, TimePostSTZ)
TimePostSTZ_metaregression


# We can build a bubble plot to visualise it

regplot(
  TimePostSTZ_metaregression, # enter your meta-regression input
  refline = 0,
  xlab = "Timing of Administration of Intervention post STZ induction (in days)",
```

```
  ylab = "SMD",
  col="tomato4",
  bg = "tomato1")

# STZ dose

# Example of renaming for simplicity
names(Nested_all_1)[names(Nested_all_1) == "STZ dose for diabetes induction"] <-
"STZdose"

Nested_all_1_MA <- metagen(
  `Nested_ES`,
  `Nested_ES_SE`,
  sm = "SMD",
  prediction = TRUE,
  data = Nested_all_1,
  studlab = StudyId,
  random = TRUE,
  fixed = FALSE,
  method.tau = "REML",
)

Nested_all_1_MA

STZdose_metaregression <- metareg(Nested_all_1_MA, STZdose)
STZdose_metaregression

regplot(
  STZdose_metaregression, # enter your meta-regression input
  refline = 0,
  xlab = "STZ dose for diabetes induction (mg/kg)",
  ylab = "SMD",
  col="tomato4",
  bg = "tomato1")



# Treatment Duration



# Example of renaming for simplicity
names(Nested_all_1)[names(Nested_all_1) == "Duration of Treatment (in weeks)_FALSE"]
<- "TreatmentDuration"

Nested_all_1_MA <- metagen(
  `Nested_ES`,
```

```
  `Nested_ES_SE`,
  sm = "SMD",
  prediction = TRUE,
  data = Nested_all_1,
  studlab = StudyId,
  random = TRUE,
  fixed = FALSE,
  method.tau = "REML",
)

Nested_all_1_MA

TreatmentDuration_metaregression <- metareg(Nested_all_1_MA, TreatmentDuration)
TreatmentDuration_metaregression

regplot(
  TreatmentDuration_metaregression, # enter your meta-regression input
  refline = 0,
  xlab = "Duration of Treatment (in weeks)",
  ylab = "SMD",
  col="tomato4",
  bg = "tomato1")
```
