## Supplementary material for "Machine-assisted systematic review and meta-analysis of citrus flavonoids in streptozotocin-induced painful diabetic neuropathy": Regular expression codes

### Regular Expression Codes NP-SOLES

<table>
<tr><th>Model</th><th>Intervention</th><th>Outcome</th></tr>
<tr>
<td>(streptozotocin|STZ|streptozocin)</td>
<td>citrus\s?flavonoid(s)?|2′-Hydroxy-3,4,4′-trimethoxy-5′,6′-\(2″,2″-dimethylpyrano\) chalcone|2′-hydroxy-3,4,4′,5-tetramethoxy-5′,6′-\(2″2″imethylpyran\) chalcone|2′-hydroxy-3,4,4′,5,6′-Pentamethoxychalcone|2′-Hydroxy-3,4,5,4′,6′-pentamethoxychalcone|2′-Hydroxy4,4′-dimethoxy-5′,6′-\(2″,2″-dimethylpyrano\) chalcone|2-\(S\)-5,7,3′,4′,5′-Pentamethoxyflavanone|3,3′,4′,5,7-Pentamethoxyflavone|3′,4′-Methylenedioxy-5,7dimethoxyflavone|3′,4′-Methylenedioxy-5,6,7-trimethoxyflavone|3,4,5 Trimethoxyflavone-7-O-arhamnoside|3,4,5-Trihydroxy-7-\(3′,7-dimethyocta-2,6-dienyloxy\) flavanone|3′,4′,5′,5,7-Pentamethoxyflavone|3′,4′,5,7,8-Pentamethoxyflavone, 5-hydroxy-3′,4′,6,7-tetramethoxyflavone|3′,4′,7,8-Tetramethoxy-5,6-\(2″,2″-dimethylpyrano\)-flavone|3′,5′-di-C-β-glucopyranosylphloretin</td>
<td>General Mechanical Outcomes:<br>(mechanical|tactile)<br>(withdrawal|response(s)?|responsiveness|pain|sensitivity|stimuli|stimulus|stimulation|threshold|pressure|allodynia|(hyper)?algesi[ac]|hypoalgesi[ac]|(hyper)?sensitivity|test(s|ing)?|nociceptive|nociception|induce)<br><br>Automated Von-Frey Testing:<br>(electronic|automated)<br>(von[ \-]Frey|semmes[ \-]weinstein)|dynamic[ \-]plantar[ \-]aesthesiometer|anesthesiometer<br><br>Manual Monofilament/Von-Frey Testing:<br>von[ \-]Frey|von[ \-]Frey(mono)?filament(s)?|von[ \-]Frey<br>hair(s)?|semmes[ \-]weinstein(hair(s)?|(mono)?filament(s)?)|force application<br><br>Pressure Pain Testing:<br>Randall[ \-]Selitto|(paw|tail)[ \-]pressure|[Aa]nalgesiometer|[Aa]nalgesimeter|[Aa]nalgesy[ \-][Mm]eter|[Aa]nalgesymeter|tail[ \-]pinch|Haffner'stest<br><br>Outcome - thermal: general:<br>\b((thermal|heat|cold)\s?(withdrawal\s?latency|pain\s?sensitivity|nociceptive\s?(threshold|test(s)?)|vocalisation\s?threshold|(hyper)?sensitivity|allodynia|hyperalgesia|stimulus|stimuli|pain|response|source|nociception|stimulation\s?(system)?))\b<br><br>Outcome - thermal – cold:<br>\b(cold\s?(withdrawal\s?latency|pain\s?sensitivity|nociceptive\s?(threshold|test(s)?)|vocalisation\s?threshold|(hyper)?sensitivity|allodynia|hyperalgesia|stimulus|stimuli|pain|response))\b<br><br>Outcome - thermal: heat:<br>\b(heat\s?(withdrawal\s?latency|pain\s?sensitivity|nociceptive\s?(threshold|test(s)?)|vocalisation\s?threshold|(hyper)?sensitivity|allodynia|hyperalgesia|stimulus|stimuli|pain|response))\b</td>
</tr>
</table>

<table>
<tr>
<td></td>
<td>|3,5-Dimethoxy-6,7;3′,4′-dimethylenedioxyflavone|3,5-Dihydroxy-7-methoxy-3,4-methylenedioxyflavone|3,5-Dihydroxy-7,8-dimethoxy-3,4-methylenedioxyflavone|3,5-Hihydroxy-7-isopentenyloxy-8-methoxy-3,4-methylenedioxyflavone|3,5,3′-Trihydroxy-8,4′-dimethoxy-7-\(3-methylbut-2-enyloxy\) flavone|3,5,4′-Trihydroxy-8,3′-dimethoxy-7-\(3-methylbut-2-enyloxy\) flavone|3,5,7-Trihydroxy-6-methoxy-4′-\(3-methylbut-2-enyloxy\) flavone|3,5,7-Trihydroxy-8-methoxy-4-\(3-methylbut-2-enyloxy\) flavone|3,5,6,7,8,3′,4′-Heptamethoxyflavone|3,5,7,3′,4′,5′-Hexamethoxyflavone|3,5,7,8,3′,4′-Hexamethoxyflavone|3,5,7,8,3′,4′,5′-Heptamethoxyflavone|3,5,8-Trimethoxy-3′,4′-methylenedioxy-7-prenyloxyflavone|3,5,8-Trimethoxy-3′,4′,6,7- bis methylenedioxyflavone|3,5,8-Trimethoxy-6,7;3′,4′-dimethylenedioxyflavone|4′-Hydroxy-3,5,3′-trimethoxy-7-prenyloxyflavone|4′-Hydroxy-3,5,7,3′-</td>
<td>Outcome - thermal: hot plate:<br>\b(hot\s?-?\s?plate\s?(test|response|latency|method)?)\b<br><br>Outcome - thermal: tail flick:<br>\b(tail\s?-?\s?(flick|withdrawal)\s?(test|latency|response|reaction)?)\b<br>Outcome - thermal: Hargreaves:<br>\b((Hargreaves|plantar)\s?(test|procedure|method)?)\b<br><br>Outcome - thermal: cold plate:<br>\b(cold\s?plate\s?(test|response|latency)?)\b<br><br>Outcome - thermal: acetone evaporation<br>\b(acetone\s?(evaporation\s?|drop\s?|spray\s?)?(test|response|assay|method)?)\b<br><br>Outcome - thermal: cold plantar assay:<br>\b(cold\s?plantar\s?(assay|test|response|latency)?)\b<br><br>Outcome - thermal: tail immersion:<br>\b((tail\s?-?\s?immersion|warm\s?-?\s?water)\s?(test|procedure|method|response)?)\b</td>
</tr>
</table>

<table>
<tr><td></td><td>tetramethoxyflavone|4,5-Dihydroxy-3,6,3′-trimethoxy-7-\(3-methylbut-2-enyloxy\) flavone|4,5-Dihydroxy-3,8,3′-trimethoxy-7-\(3-methylbut-2-enyloxy\) flavone|4′,5,7-Trihydroxy-3,3′,8-trimethoxyflavone|5-Demethylnobiletin|5-Hydroxy-3-isopentenyloxy-7-methoxy-3,4-methylenedioxyflavone|5-Hydroxy-3,8-dimethoxy-6,7;3′,4′-dimethylenedioxyflavone|5 O-desmethylnobiletin|5,4′-Dihydroxy-3,6,7,8,3′ pentamethoxyflavones|5,4′-Dihydroxy-3,7,3′-trimethoxyflavone|5,4-Dihydroxy-3,7,8,3′-tetramethoxy flavone|5,6,3′-Trihydroxy-7,4′-dimethoxyflavone|5,6,7,3′,4′,5′-Hexamethoxyflavone|5,6,7,3′,4′,5′-Hexamethoxyflavanone|5,7-Dihydroxy-3-methoxy-4′-\(3-methylbut-2-enyloxy\) flavone|5,7-Dihydroxy-3,3′,8-trimethoxy-4′-\(3-methylbut-2-enyloxy\) flavone|5,7-Dihydroxy-3,6-dimethoxy-4-\(3′-methylbut-2-enyloxy\) flavone|5,7-Dihydroxy-3,8-dimethoxy-4′-\(3-methylbut-2-enyloxy\) flavone|5,7-Dihydroxy-8-methoxy-3,4-di\(3-</td><td></td></tr>
</table>

| | |
|---|---|
| | methylbut-2-enyloxy\) flavone\|5,7,3′,4′,5′-Pentamethoxyflavone\|5,7,3′,4′,5′-Pentamethoxyflavanone\|5,7,8,3′,4′,5′-Hexamethoxyflavone\|6-Farnesyl-5,7,3′,4′-tetrahydroxy flavanone\|7-Methoxyflavone\|acacetin 3,6-di-C-glucoside\|apigenin\|apigenin 8-C-neohesperidoside\|auranetin\|daidzin\|dihydrokaempferol\|dihydrokaempferol 3-O-rhamnoside\|diosmin\|diosmin 1\|engeletin\|evodioside-B\|excavasideA\|excavasideB\|flavaprin\|furanocoumarin \[5-methoxy-8-geranyloxypsoralen]\|genistin\|glychalcone\|hesperidin\|hesperitin\|hesperitin-7-O-β-D-glucopyranoside\|hispidulin\|homoplantaginin\|icariside\|isokanugin\|isonaringin\|isoquercitrin\|kaempferide-3-O-β-D-glucopyranoside\|kaempferol\|kaempferol-3-O-β-D-glucopyranoside\|kaempferol 3-O-β-D-glucopyranosyl \(1–2\)-α-D-xylopyranoside\|kaempferol-3-O-rutinoside\|kaempferol 3-robinobioside\|kumatakenin \(5,4′-dihydroxy-3,7-dirnethoxyflavone\) 9avone\|lemairone\|lucen |

<table>
<tr><td></td><td>in-2 4′-methyl ether|luteolin|luteolin 8-C-β-D-glucopyranoside|marmesin|melibentin|melibentin \[3,5,6,7,8-pentamethoxy-3′,4′-methylenedioxyflavone]|meliflavonesA|meliflavonesB|melisimplexin|melisimplexin \[3,5,6,7-tetramethoxy-3′,4′-methylenedioxyflavone]|meliternatin|myricetin 3′-α-l-rhamnopyranoside|myricetin 3,3′-di-α-l-rhamnopyranoside|myricetin 3-O-β-D-rutinoside|naringenin|naringenin-7-O-β-D-glucopyranoside|naringin|narirutin|narirutin 4′-O-glucoside|neoeriocitrin|neohesperidin|neoponcirin|nepitrin|nobiletin|phloretin 3′,5′-di-C-glucoside|poncirin|quercetin|rhamnetin|rhoifolin|rutin|sinensetin|tamarixetin 3-robinobioside|tangeritin|vicenin-2|vitexin</td><td></td></tr>
</table>

*Table 1. Regular expression codes for streptozotocin induced painful diabetic neuropathy animal models, citrus flavonoid treatments and mechanical-pain related outcomes*
